# PARP1 Exhibits an Enzymatically Inactive Chromatin Binding Mode

**DOI:** 10.64898/2026.06.24.734344

**Authors:** Alexandria C. Fiorenza, Mahika Anand, Karolin Luger

## Abstract

Poly (ADP-ribose) Polymerase 1 (PARP1) is an abundant nuclear enzyme that dynamically engages chromatin in diverse cellular scenarios. In the context of DNA repair, PARP1 becomes enzymatically activated and subsequently attaches ADP-ribose units onto various proteins, including histones, to signal and coordinate the DNA damage response. In the absence of DNA damage, PARP1 modulates chromatin structure by directly binding to nucleosomes, however, the molecular basis of this interaction is unknown. Here, we define a distinct, enzymatically inactive mode of PARP1 chromatin binding, in which the Zn1, Zn2, Zn3, and BRCT domains cooperatively bind nucleosomal linker DNA and drive compaction of undamaged chromatin. This binding mode does not trigger catalytic activation and therefore is insensitive to PARP inhibitors (PARPi). Together, our results support a model in which PARP1 associates with the genome in an inactive state to compact chromatin and to surveil for DNA lesions.

**Summary:** PARP1 engages undamaged chromatin in a distinct binding mode that results in chromatin compaction but does not lead to enzymatic activation.

## INTRODUCTION

Chromatin is critical for condensing DNA, maintaining genome integrity, and regulating gene expression. The fundamental building block of eukaryotic chromatin is the nucleosome, a nucleoprotein complex containing ∼147 bp of DNA wrapped around the histone octamer complex (*1*). Nucleosomes are connected by “linker” DNA (regions of DNA ranging from ∼20-100 bp) to form nucleosome arrays in a “beads-on-a-string” arrangement (*2–4*). Epigenetic modifications to the canonical nucleosome, such as incorporation of histone variants and histone tail modifications, as well as linker histones and chromatin architectural proteins, alter nucleosome stability and structure, further promoting compact or relaxed chromatin states (*5–8*). Different chromatin states regulate DNA accessibility, which in turn controls any nuclear process that requires access to DNA. New insights into eukaryotic chromatin architecture continue to emerge as more proteins are implicated in shaping, moving, or altering nucleosomes during various cellular processes (*9*, *10*). Understanding how different proteins drive distinct chromatin states is essential in unraveling the mechanisms by which the eukaryotic genome is structured and organized.

Poly (ADP-ribose) Polymerase 1 (PARP1) is a highly abundant, chromatin-associated enzyme that alters nucleosome stability and structure (*11–13*). PARP1 is widely known for its role in the DNA damage response, in which PARP1 detects and signals the presence of many different types of DNA lesions (*12*, *14*). Upon binding to damaged DNA, PARP1 undergoes a conformational change that uncovers the active site (*15*). This catalytically activated form catalyzes the attachment of poly-ADP-ribose (PAR) units from NAD^+^ onto itself in a process termed auto-PARylation (*16*, *17*). Activated PARP1 also forms a composite active site with Histone PARylation Factor 1 (HPF1) which enables PARylation of histones tails (*18–20*). Histone PARylation is responsible for destabilizing the nucleosome and ‘relaxing’ chromatin to allow PAR-recruited chromatin remodelers and the DNA repair machinery to access the damage site (*21–23*). Historically, PARP1 has been studied in this context due to the synthetic lethal nature of PARP1 inhibition in DNA repair deficient cancers (*24–26*). Hence, various PARP inhibitors (PARPi) have been developed and approved for clinical use that efficiently kill cancer cells lacking proficient BRCA-mediated DNA repair (*27*, *28*). However, PARP1 has been implicated in many other cellular processes (i.e. transcription, immune response, cell death, etc.) that are much less understood (*13*, *29*, *30*). And further, despite their clinical success, it is unknown whether PARPi disrupt these alternate PARP1 functions in healthy cells.

In the absence of DNA damage, PARP1 contributes to the regulation of gene expression in ways that are not well-understood (*13*, *31–34*). For instance, PARP1 associates with both active and inactive epigenetic marks, binds various regulatory DNA elements, stabilizes DNA loops, and directly alters chromatin structure through nucleosome compaction (*35–39*). Functionally, these mechanisms suggest a strong interaction between PARP1 and undamaged DNA/chromatin. Indeed, in vitro binding assays validate that PARP1 binds both plasmid DNA and nucleosome arrays with high affinity (*40*, *41*). Furthermore, PARP1 induces chromatin compaction by directly binding nucleosomes, resulting in nuclease resistant complexes that repress transcription (*36*, *42*). This function in particular is independent of its catalytic activity; however, PARP1 activation is intimately coupled to its interaction with DNA. As such, how it engages undamaged chromatin and DNA while remaining inactive is unknown.

PARP1 is a large enzyme that consists of six folded domains: three zinc fingers (Zn1, Zn2, Zn3), a BRCA 1 C-terminal domain (BRCT), a tryptophan-glycine-arginine (WGR) rich-motif, and a catalytic (CAT) domain that contains a helical (HD) and ADP-ribosyl-transferase (ART) subdomain (**Fig. 1A**). Structural and biophysical characterization of PARP1 reveal that the Zn1, Zn2, Zn3, and WGR domains engage DNA lesions by binding the exposed nitrogen bases and DNA backbone (*43–47*). However, the lack of exposed nitrogen bases in undamaged chromatin infers an alternate PARP1-chromatin binding mode that is incompatible with the reported DNA damage binding modes. The molecular basis of how PARP1 engages and compacts chromatin in the absence of DNA damage remains largely unexplored, limiting our fundamental understanding of its biology. Furthermore, characterizing its different binding modes could contribute to the development of more specific inhibitors, as it is unknown whether current PARPi disrupt these lesser studied functions. In this study, we biochemically characterize the chromatin architectural binding mode of PARP1. We found that PARP1 compacts chromatin in a distinct binding mode that does not lead to catalytic activation. Specifically, the three zinc fingers and BRCT domains engage the DNA linkers of undamaged chromatin. In this conformation, the HD does not unfold, and PARP1 remains inactive. Additionally, we found that both clinically approved and in-trial PARPi do not alter the interaction of PARP1 with undamaged chromatin.

**Figure 1.**
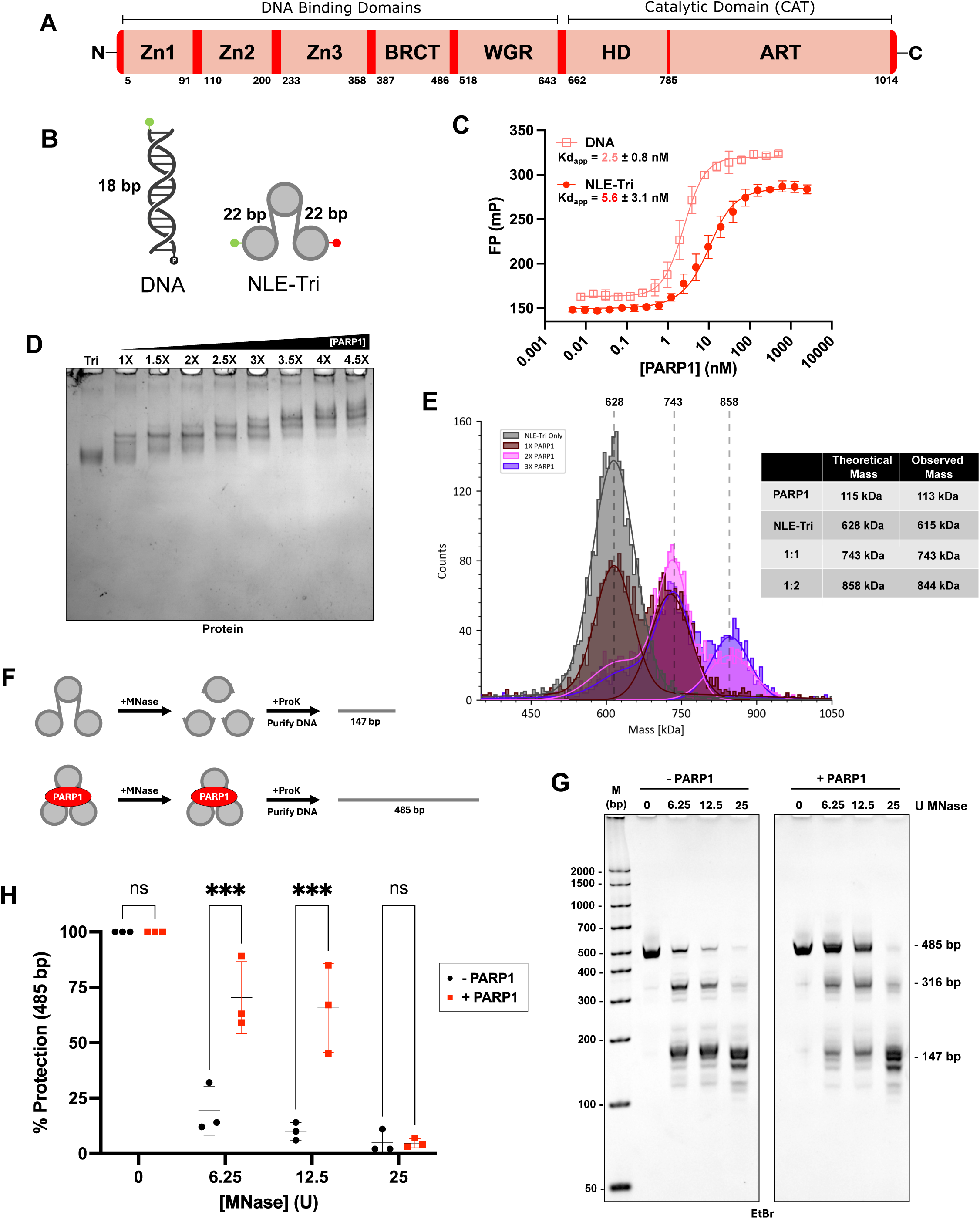
PARP1 binds nucleosomal linker DNA with high affinity. (**A**) Schematic of PARP1 domains. (**B**) DNA and chromatin substrates: p18mer DNA and NLE-Tri-22 with Alexa 488 fluorophore (green), Alexa 647 fluorophore (red), and phosphate (black) attached to the 5’ ends of DNA. (**C**) Fluorescence polarization (FP) binding curves of PARP1 bound to p18mer DNA (pink square) and NLE-Tri-22 (red circle). Apparent dissociation constants (Kd_app_) with standard deviations over n=4 replicates are given. The curve obtained for p18mer was fit to the four-parameter logistic curve whereas NLE-Tri was fit to the quadratic equation (see methods). (**D**) EMSA of PARP1-NLE-Tri-22 complex with increasing concentrations of PARP1 titrated into a fixed concentration of NLE-Tri (50 nM). PARP1 molar equivalences are indicated above each lane. (**E**) Mass photometry analysis of PARP1-NLE-Tri-22 complexes with increasing concentrations of PARP1 compared to NLE-Tri alone (grey). Theoretical molecular weights for each complex are indicated on the histogram in grey dashed lines and provided in the table. One representative observed mass is reported in the table, all individual masses are reported in Fig. S2. (**F**) Schematic for the MNase protection assay in the presence and absence of PARP1 (**G**) Representative MNase digest, analyzed by PAGE. Trinucleosomes (200 nM) in the presence (400 nM) and absence of PARP1 with increasing units of MNase, as indicated above each lane. (**H**) Quantitative analysis of the percent protection of the 485 bp band with increasing units of MNase. FIJI was used to determine each 485 bp band intensity. Each 485 bp band intensity was normalized to the corresponding 485 bp control (0 U) in the presence (red square) and absence (black circle) of PARP1 to determine the percent protection. Statistical significance was determined using the unpaired t-test with Bonferroni-Dunn multiple comparison correction p<0.001 (***), n=3.

## RESULTS

### PARP1 binds nucleosomal linker DNA with high affinity

Since the 1970’s, PARP1 has been proposed to bind chromatin linker DNA in a manner analogous to histone H1 (*40*, *42*, *48–52*). In vitro characterizations of PARP1 bound to various nucleosome constructs suggest that PARP1 engages the DNA linkers extending from the nucleosome rather than the nucleosome core. However, these nucleosome constructs contain exposed DNA ends that mimic double stranded DNA breaks, a preferred substrate for PARP1 binding and activation. Previously, PARP1 has been shown to bind a ‘non-linker ended’ (NLE) trinucleosome, where three nucleosomes are separated by 60 bp of linker DNA with no additional flanking DNA (termed NLE-Tri-60). Because NLE-Tri-60 lacks accessible DNA ends, it is an appropriate minimalist model for undamaged chromatin. In our previous study, we showed that one PARP1 molecule bound NLE-Tri-60 with high affinity (*40*), further suggesting a strong interaction with the internal linkers.

In eukaryotic cells, chromatin linkers can range from ∼20-100 bp (*53*, *54*). Therefore, we wanted to test whether chromatin linkers shorter than 60 bp altered PARP1 engagement. Here, we chose to use a biologically relevant short linker length of 22 bp. To do this, we developed NLE-Tri-22, a dual-labeled NLE trinucleosome with 22 bp linkers between nucleosomes and no accessible DNA ends (**Fig. 1B, S1A-G**). NLE-Tri-22 was designed with three different Widom positioning sequences and PCR amplified to attach AlexaFluor 488 and AlexaFluor 647 on either 5’ end of the DNA (**Fig. S1A-C**). Fluorescence polarization was first employed to determine the binding affinity of PARP1 to NLE-Tri-22 compared to short linear DNA (18mer) (**Fig. 1B**). We found that PARP1 binds NLE-Tri-22 with high affinity: 5.6 nM, compared to the 18mer DNA to which it binds with an affinity of 2.5 nM (**Fig. 1C**, **Table 1**). PARP1 affinity for NLE-Tri-22 was similar to the reported affinity for NLE-Tri-60 (4.8 nM (*40*)), indicating that shorter linkers do not affect PARP1 binding.

**Table 1.** PARP1 binding affinities at 100 mM KCl. Reported apparent dissociation constants (Kd_app_) for full-length (FL) PARP1 and single domain deletion constructs bound to 18mer DNA and NLE-trinucleosomes at 100 mM KCl. The affinities obtained for p18mer were determined from the four-parameter logistic curve whereas NLE-Tri affinities were determined from the quadratic equation. Hill coefficients, R^2^, and replicate values are reported.

| Construct | DNA (p18mer) |  |  |  | Undamaged Chromatin (NLE-Tri) |  |  |
| --- | --- | --- | --- | --- | --- | --- | --- |
| | $K_{d_{app}}$ | Hill Coeff. | $R^2$ | Replicates | $K_{d_{app}}$ | $R^2$ | Replicates |
| FL | $2.5 \pm 0.8$ | 1.69 | 0.951 | n=4 | $5.6 \pm 3.1$ | 0.986 | n=4 |
| $\Delta Z_{n1}$ | $2.3 \pm 0.2$ | 2.09 | 0.974 | n=4 | $10.1 \pm 3.6$ | 0.987 | n=4 |
| $\Delta Z_{n2}$ | $2.8 \pm 1.3$ | 1.82 | 0.912 | n=4 | $8.2 \pm 1.8$ | 0.996 | n=4 |
| $\Delta Z_{n3}$ | $2.4 \pm 1.5$ | 1.53 | 0.951 | n=4 | $3.3 \pm 1.2$ | 0.986 | n=4 |
| $\Delta BRCT$ | $4.6 \pm 1.0$ | 1.85 | 0.979 | n=4 | $66.7 \pm 22.3$ | 0.993 | n=4 |
| $\Delta WGR$ | $1.2 \pm 0.4$ | 1.28 | 0.939 | n=4 | $10.6 \pm 3.0$ | 0.991 | n=4 |
| $\Delta CAT$ | $5.6 \pm 2.3$ | 1.76 | 0.950 | n=4 | $12.7 \pm 6.1$ | 0.990 | n=4 |
| N-PARP | | | | | $10.4 \pm 6.2$ | 0.944 | n=3 |

Because the affinities between both undamaged chromatin constructs (NLE-Tri-60 and NLE-Tri-22) are similar, we hypothesized that one PARP1 molecule should also bind to the NLE-Tri-22 linkers extending from the central nucleosome, as suggested for NLE-Tri-60. We used Electrophoretic Mobility Shift Assay (EMSA) and Mass Photometry (MP) to probe the stoichiometry of the PARP1-NLE-Tri-22 complex. EMSA analysis revealed several shifted bands in the presence of increasing amounts of PARP1, suggesting that more than one PARP1 molecule binds NLE-Tri-22 (**Fig. 1D**). We analyzed the same samples using Mass Photometry (MP), which enables the determination of the molecular weight of the complex in solution under equilibrium conditions. This analysis reveals two distinct bound complex populations at ∼743 and ∼858 kDa, corresponding to NLE-tri-22 bound to one and two PARP1 molecules, respectively (**Fig. 1E, S2C-G**). Even in the presence of excess PARP1, a higher percentage of 1:1 complex was seen compared to the 1:2 complex, suggesting the preferential formation of 1:1 complex (**Fig. S2E-G**). PARP1 concentrations exceeding a three-fold excess resulted in precipitation that could not be analyzed by MP.

We next sought to determine if PARP1 was bound to the linker DNA, as previously proposed for the NLE-tri-60 substrate. To assess this, we used a Micrococcal Nuclease (MNase) protection assay. In this assay, NLE-Tri-22 was digested with increasing units of MNase in the presence and absence of PARP1 (**Fig. 1F**). Samples were then treated with Proteinase K (ProK) to remove histones and bound PARP1 to allow analysis of the resulting DNA fragment sizes. As the nucleosome protects ∼147 bp of DNA, protection of linker DNA by PARP1 would result in DNA fragments larger than 147 bp; complete protection would result in the full-length DNA fragment of 485 bp. We decided to add a two-fold excess PARP1 over NLE-tri-22, which primarily contained a 1:1 complex, as shown by EMSA and MP analysis (**Fig. 1D & E**). In the presence of PARP1, we observed more full-length 485 bp DNA fragment, while in the absence of PARP1, DNA was digested readily to ∼147 bp length (**Fig. 1G**). To quantify the percent protection, the intensity of each 485 bp band was normalized to its corresponding control (0 U MNase). There was significantly more protection of the 485 bp band in the presence of PARP1 (**Fig. 1H**), demonstrating that PARP1 is bound to linker DNA. As the full 485 bp DNA was protected in the presence of one molecule of PARP1, this also suggests that one PARP1 molecule can bind both linkers simultaneously, consistent with previous studies suggesting that PARP1 bridges two DNA molecules (*35*, *41*). Altogether, we have found that PARP1 binds the linker region of NLE-Tri-22 with high affinity. Our results suggest that PARP1 can access chromatin linker DNA as short as ∼20 bp in length, consistent with the notion that PARP1 protects ∼10 bp of DNA (*42*). Hereafter, NLE-Tri-22 will be termed NLE-Tri for simplicity.

### PARP1 compacts trinucleosomes by binding to the connecting linker DNA

Binding of PARP1 to nucleosome arrays promotes compacted chromatin structures, which entails bringing individual nucleosomes into closer proximity with each other (*36*, *40*). Previous studies exploring PARP1-induced chromatin compaction typically utilize methods that rely on imaging (i.e. atomic force microscopy), which is limited to mostly qualitative analysis. To quantitatively assess chromatin compaction, we developed a Förster Resonance Energy Transfer (FRET) assay using NLE-Tri in which the DNA ends are labeled with fluorescence donor and acceptor, respectively. In absence of compaction, there should be little to no FRET signal as the fluorophores are not within the Förster distance (**Fig. 2A**). When nucleosomes are brought into closer proximity, excitation at the 488 nm wavelength will produce increased emission at the 647 nm wavelength. The intensity of the 647 nm wavelength following 488 excitation (FRET intensity) can be normalized to the intensity of the 488 nm wavelength following 488 excitation (Donor intensity) to determine FRET ratio. Upon titration of PARP1, we observed an increase in FRET ratio (**Fig. 2B**). Due to the propensity of PARP1-NLE-Tri complexes to aggregate at excess PARP1 concentrations, as seen in MP, the FRET signal becomes noisy beyond 60 nM PARP1. We therefore selected 50 nM PARP1 (equivalent to ∼2.5X) to proceed with replicates (**Fig. 2B, black arrow**). In the presence of PARP1, there was a significantly increased FRET ratio compared to the absence of PARP1 (**Fig. 2C**), suggesting that PARP1 compacts NLE-Tri.

**Figure 2.**
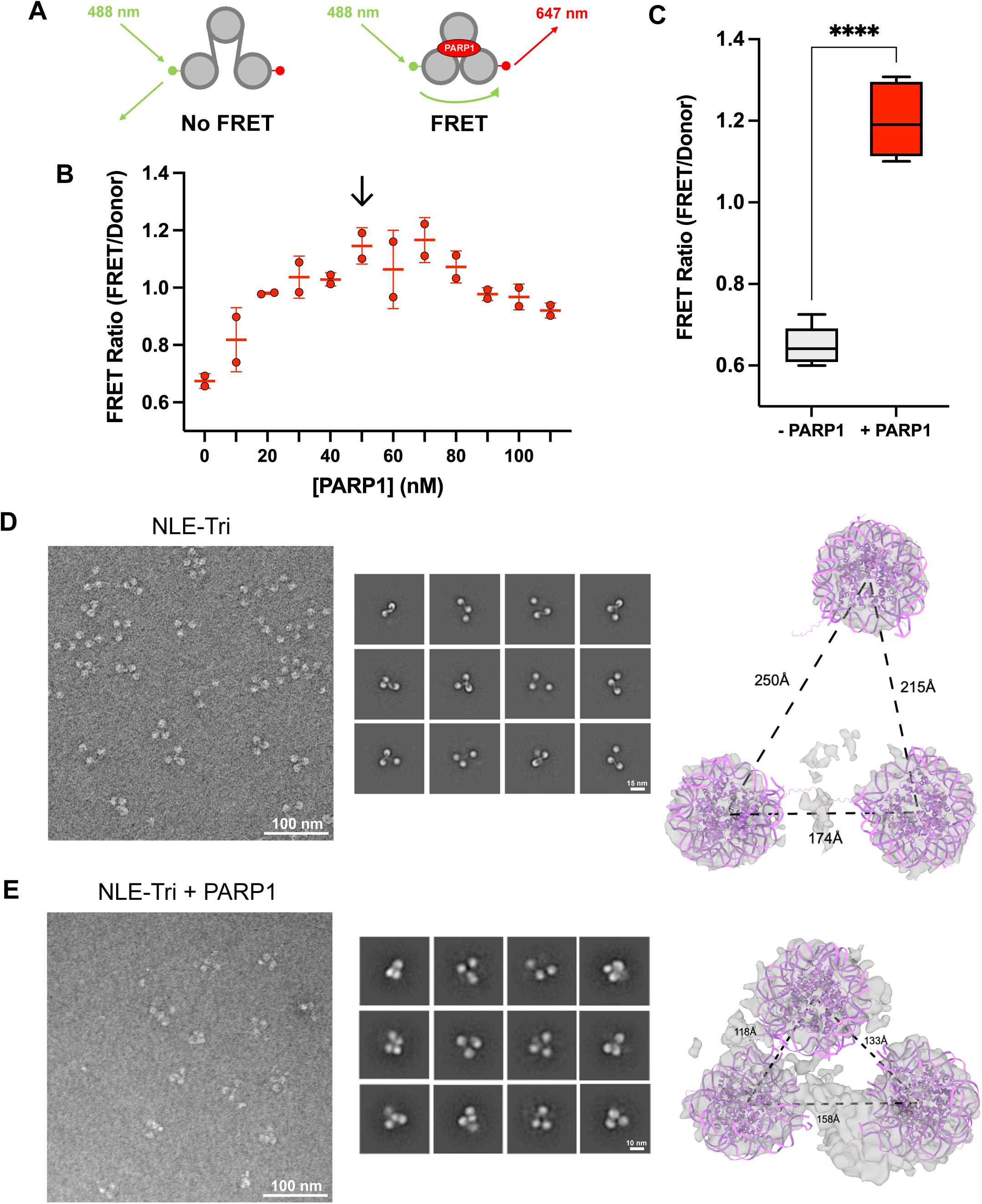
PARP1 compacts NLE-Tri. (**A**) Schematic for FRET-based chromatin compaction assay. AlexaFluor 488 (green) and AlexaFluor 647 (red) fluorophores are attached to 5’ ends of the 485 bp DNA. (**B**) FRET ratio is plotted for a titration range of PARP1, as indicated, to 20 nM NLE-Tri. FRET ratio was determined by normalizing each FRET intensity to the 488 nm donor intensity. Addition of excess PARP1 (beyond a ratio of 60 nM PARP1 to 20 nM NLE-Tri) leads to a decrease in FRET ratio, likely due to quenching by aggregation. 50 nM PARP1 (2.5X molar equivalents over NLE-Tri) was chosen for replicates (black arrow). (**C**) FRET replicates at 50 nM PARP1 and 20 nM NLE-Tri. Statistical significance was determined using the unpaired t-test p<0.0001 (****); n=5. (**D; E**) Assessment of PARP1 interaction without **(D)** and with NLE-Tri (**E)** by negative stain EM. Representative micrograph (left, scale bar=100 nm), 2D classification (center, scale bar=15 nm (D), scale bar=10 nm (E)), and *ab initio* reconstruction (right, grey density). Mononucleosome (pink, PDB: 1AOI) was fit into each nucleosome-like density. Distances between nucleosomes were measured in ChimeraX by defining the centroid of each mononucleosome.

To validate the compaction observed via FRET assay, we analyzed the 2.5X complex by negative stain TEM and compared it to NLE-Tri in absence of PARP1 (**Fig. 2D & E**). CryoSPARC was used to process both datasets, resulting in ab initio reconstructions of the averaged trinucleosomes (**Fig. S3**). Using ChimeraX, a mono-nucleosome (PDB: 1AOI) was fit into each nucleosome-like density. The centroids of each nucleosome were defined. In absence of PARP1, the centroid distances between the three nucleosomes were 250 Å, 215 Å, and 174 Å (**Fig. 2D**). In the presence of PARP1, the centroid distances decreased to approximately 118 Å, 133 Å, and 158 Å (**Fig. 2E**). Additional density was observed in the presence of PARP1 (**Fig. 2E, extra grey density**), which we believe to be PARP1. However, due to low resolution of negative stain TEM, we are unable to confidently determine its orientation.

### N-terminal PARP1 is Required for PARP1-Induced Chromatin Compaction

Our results show that PARP1 induces compaction of NLE-Tri, demonstrating that PARP1 can compact undamaged chromatin by binding to biologically relevant connecting linker DNA. However, which domains of PARP1 engage the chromatin linker is unknown. In the context of double stranded or single stranded DNA breaks, PARP1 binds the exposed nitrogen bases and the DNA backbone through engagement of the three zinc fingers and WGR domain (*43–47*). Since undamaged chromatin lacks exposed nitrogen bases, we expect PARP1 to engage undamaged DNA in an alternate conformation, using different domains. To determine which domains interact with NLE-Tri we tested the ability of previously established PARP1 single domain deletion constructs (*55*, *56*) to bind and compact NLE-Tri (**Fig. 3A, S4A**). EMSA was first employed to show that each of the domain deletions retained the ability to bind NLE-Tri (**Fig. S4B-G**). However, ΔBRCT (**Fig. S4E**) required more protein to shift the band compared to full-length PARP1(FL) (**Fig. 1D**), suggesting a mild disruption in binding when the BRCT domain is deleted. To more quantitatively compare affinities, fluorescence polarization was employed for each domain deletion and compared to FL PARP1. We first tested binding at 100 mM KCl and found that ΔBRCT bound NLE-Tri with decreased affinity (66.7 nM) compared to FL (5.6 nM), (**Fig. S5A, Table 1**). Due to detection limitations and the tight binding of PARP1, we hypothesized that the contribution of other PARP1 domains might not be detectable. We therefore repeated these measurements at 200 mM KCl to modulate electrostatics, revealing reduced affinities for both ΔZn2 and ΔBRCT (**Fig 3B**, **Table 2**). Neither of these domains appear to contribute to DNA binding at 100 mM or 200 mM KCl (**Fig 3C, S5B, Table 1**, **Table 2**), suggesting that Zn2 and BRCT contribute specifically to the interaction with undamaged chromatin.

**Figure. 3.**
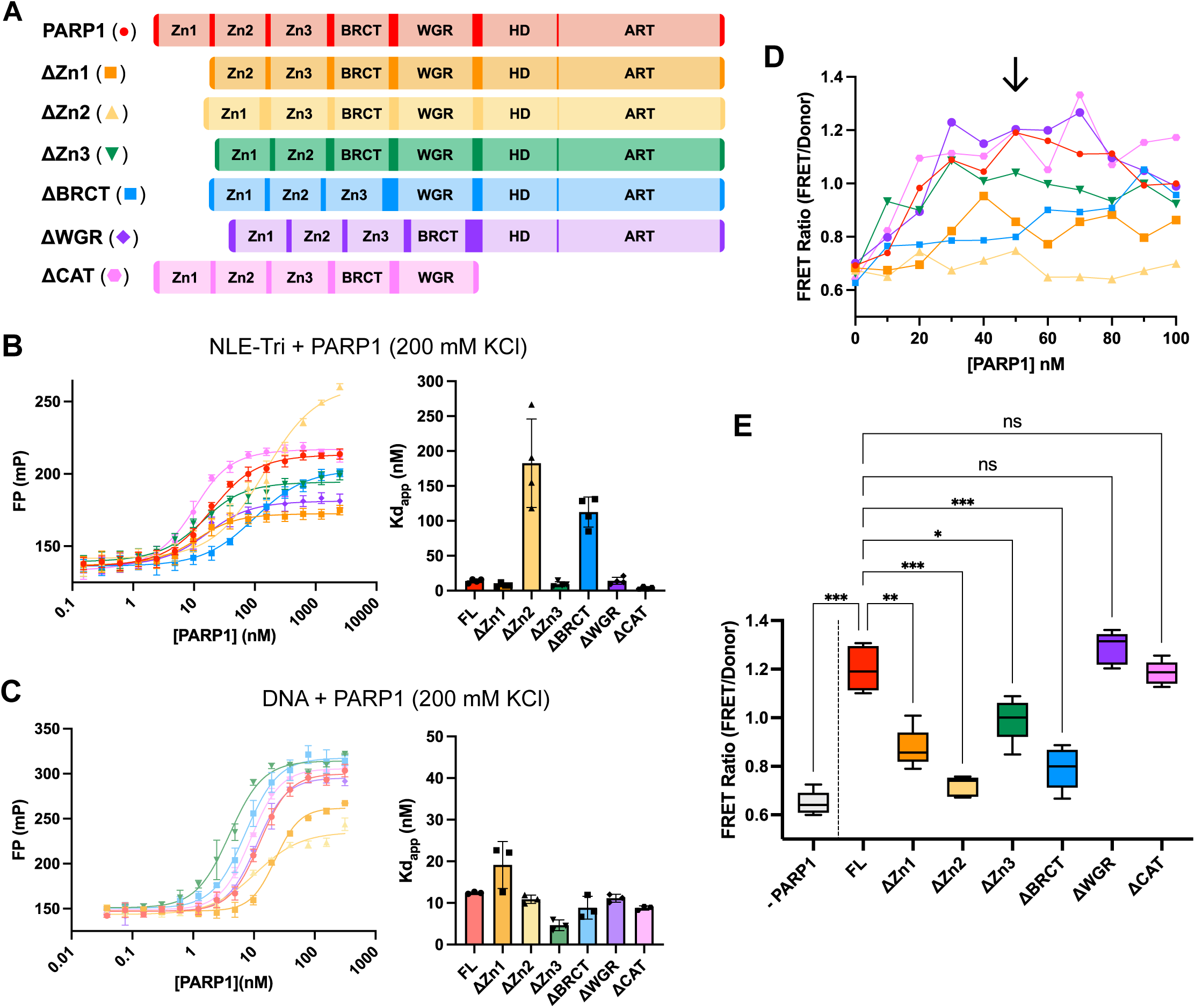
Zn1, Zn2, Zn3, and BRCT domains contribute to PARP1-chromatin binding and compaction. (**A**) Schematic of domain composition of full length PARP1 and single domain deletions. Symbols and colors shown to the left are used for all other figure panels. All constructs except ΔZn1 and ΔCAT have the deleted domain replaced with a 30 aa Ala-Gly-Leu-Ser) linker. (**B**) Fluorescence polarization curve fit to the quadratic equation (left), and replicate affinities (right) of each construct bound to NLE-Tri at 200 mM KCl, n=4. (**C**) Fluorescence polarization curve fit to the four-parameter logistic curve (left) and replicate affinities (right) of each construct bound to p18mer at 200 mM KCl, n=3. (**D**) FRET-based chromatin compaction assay of PARP1/domain deletions with NLE-Tri. FRET ratio is plotted over a titration of constructs. FRET ratio was determined by normalizing each FRET intensity to the 488 nm donor intensity. 50 nM protein was chosen for replicates (black arrow). (**E**) FRET replicates at 50 nM protein, n=5. Statistical significance was determined using the Brown-Forsythe and Welch ANOVA with Dunnett’s T3 multiple comparisons test p<0.0332 (*), p<0.0021 (**), p<0.0002(***), p<0.0001(****).

**Table 2.**
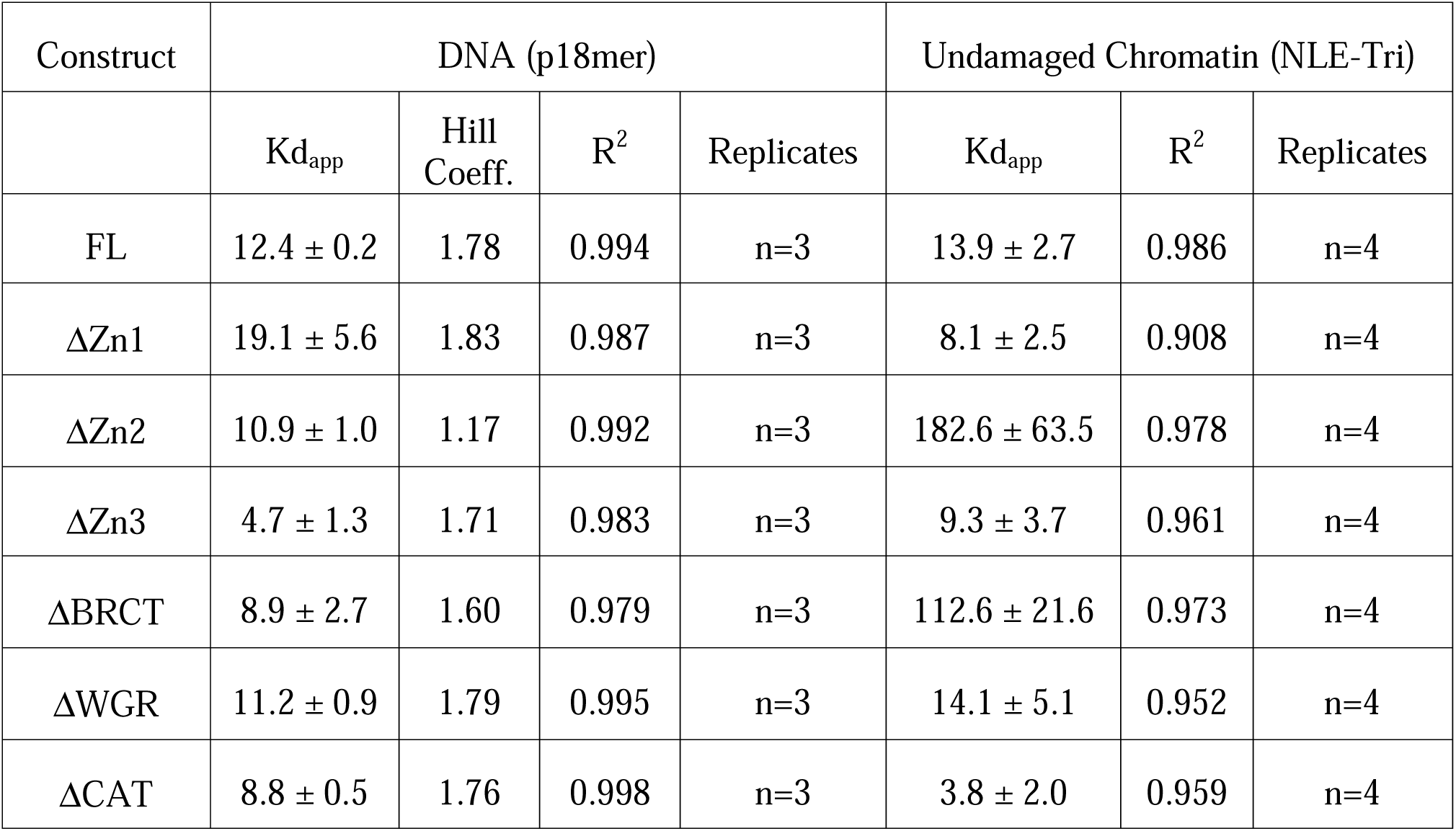
PARP1 binding affinities at 200 mM KCl. Reported apparent dissociation constants (Kd_app_) for full-length (FL) PARP1 and single domain deletion constructs bound to 18mer DNA and NLE-trinucleosomes at 200 mM KCl. The affinities obtained for p18mer were determined from the four-parameter logistic curve whereas NLE-Tri affinities were determined from the quadratic equation. Hill coefficients, R^2^, and replicate values are reported.

| Construct | DNA (p18mer) |  |  |  | Undamaged Chromatin (NLE-Tri) |  |  |
| --- | --- | --- | --- | --- | --- | --- | --- |
| | $K_{d_{app}}$ | Hill Coeff. | $R^2$ | Replicates | $K_{d_{app}}$ | $R^2$ | Replicates |
| FL | $12.4 \pm 0.2$ | 1.78 | 0.994 | n=3 | $13.9 \pm 2.7$ | 0.986 | n=4 |
| $\Delta Zn1$ | $19.1 \pm 5.6$ | 1.83 | 0.987 | n=3 | $8.1 \pm 2.5$ | 0.908 | n=4 |
| $\Delta Zn2$ | $10.9 \pm 1.0$ | 1.17 | 0.992 | n=3 | $182.6 \pm 63.5$ | 0.978 | n=4 |
| $\Delta Zn3$ | $4.7 \pm 1.3$ | 1.71 | 0.983 | n=3 | $9.3 \pm 3.7$ | 0.961 | n=4 |
| $\Delta BRCT$ | $8.9 \pm 2.7$ | 1.60 | 0.979 | n=3 | $112.6 \pm 21.6$ | 0.973 | n=4 |
| $\Delta WGR$ | $11.2 \pm 0.9$ | 1.79 | 0.995 | n=3 | $14.1 \pm 5.1$ | 0.952 | n=4 |
| $\Delta CAT$ | $8.8 \pm 0.5$ | 1.76 | 0.998 | n=3 | $3.8 \pm 2.0$ | 0.959 | n=4 |

To complement this analysis, we tested each PARP1 domain deletion in the FRET-based chromatin compaction assay. Titrations of each domain deletion were first performed (**Fig. 3D**). Again, we chose to repeat replicates at 50 nM protein (∼2.5X) to ensure we measured compaction and not aggregation (**Fig. 3D, black arrow**). The ΔZn1, ΔZn2, ΔZn3, and ΔBRCT domain deletions showed decreased FRET ratio compared to FL, whereas the ΔWGR and ΔCAT showed no differences (**Fig. 3E**). We saw the most significant reduction in FRET ratio when either the Zn2 or BRCT domains were deleted. Interestingly, ΔZn1 and ΔZn3 also had significant reductions in FRET but were not detected in our binding assay, suggesting that Zn1 and Zn3 also contribute to chromatin binding/compaction. ΔZn3 showed only moderate reduction in FRET, which suggests that its contribution is less critical than that of Zn1, Zn2, or BRCT. From these results, we conclude that the four N-terminal PARP1 domains, Zn1, Zn2, Zn3, and BRCT, work together to bind and compact undamaged chromatin.

To confirm this, we surmised that a protein construct containing only Zn1, Zn2, Zn3, and BRCT (the N-terminal half of PARP1), should still bind and compact chromatin, while the C-terminal domain of PARP1, a protein construct containing only WGR and CAT, should not (**Fig. S6A-B**). Indeed, N-terminal PARP1 (N-PARP) was able to bind NLE-Tri with similar high affinity (11.3 nM) compared to FL, whereas C-terminal PARP1 (C-PARP) did not bind (**Fig. S6C, Table 1**). We also tested N-PARP in the FRET-based chromatin compaction assay and saw an increased FRET ratio, similar to FL (**Fig. S6D**). C-PARP was unable to produce an increased FRET ratio, confirming its inability to bind (**Fig. S6D**). To confirm N-PARP binding to NLE-Tri linker DNA, we utilized the MNase protection assay. In the presence of N-PARP, the 485 bp band persisted, compared to the control in the absence of N-PARP (**Fig. S6E**). After quantification of the 485 bp bands, we found there was a significantly higher degree of protection in the presence of N-PARP (**Fig. S6F**), suggesting N-PARP is bound to and protects the linker from MNase digestion. As was observed with full length PARP1, the entire length of the DNA was protected in the presence of one equivalent of N-PARP1, suggesting that it binds both linkers simultaneously. Altogether, our data show that the N-terminal DNA binding region of PARP1 is sufficient to bind and compact NLE-Tri, validating that Zn1, Zn2, Zn3, and BRCT are integral for this interaction, while the catalytic domain and the WGR domain do not contribute to this particular function of PARP1.

### PARP1 is Catalytically Inactive in the Presence of Undamaged Chromatin

Using NLE-Tri as a proxy for undamaged chromatin, we wanted to know whether PARP1 is indeed inactive in its chromatin binding mode. In a previous study from our lab, we showed that PARP1 is inactive on undamaged plasmid DNA (*41*), and we hypothesized that PARP1 should also be inactive on undamaged chromatin. This is supported by the notion that engagement of the WGR domain is essential for PARP1 activation, as it leads to HD unfolding which frees up the catalytic binding pocket (*15*). We report here a lack of WGR engagement in the undamaged chromatin binding mode, which also suggests that PARP1 is inactive. Because PARP1 is rapidly activated upon detecting DNA lesions, PARP1 can also be activated by minute amounts of contaminating DNA. Additionally, nucleosomal DNA has the inherent ability to “breathe”, a phenomenon where DNA ends dissociate and re-bind on the histone octamer at a rapid time scale (*57*, *58*). In the context of non-linker ended nucleosomes, this can lead to the otherwise inaccessible DNA ends becoming transiently accessible, resulting in PARP1 activation. As such, to test PARP1 activity in the presence of undamaged chromatin, we first treated the NLE-Tri samples with exonucleases to remove contaminating or partially dissociated DNA (**Fig. 4A**). PARP1 and NAD+ were then added and the samples were assayed for ADPr by western blot. Without exonuclease treatment, we saw robust auto-PARylation of PARP1, whereas no auto-PARylated PARP1 was observed after treatment with exonuclease (**Fig. 4B**). This result suggests that PARP1 is inactive on undamaged chromatin but can be activated by minute amounts of free DNA or dissociated DNA ends. To confirm that NLE-Tri was still intact following exonuclease treatment, we added ProK and purified the resulting DNA fragments. The purified NLE-Tri sample with and without exonuclease showed identical bands at 485 bp, while the DNA control was completely digested, confirming that NLE-Tri is still intact (**Fig. 4C**). To test whether PARP1 was still enzymatically active even after having been exposed to exonuclease, we quenched the exonuclease reaction, then added dsDNA. This resulted in a robust ADPr signal, showing that PARP1 is still competent for activation upon encountering DNA damage (**Fig. 4A, magenta lane**). These data demonstrate that PARP1 is indeed enzymatically silent on undamaged chromatin and that DNA contamination can result in inadvertent activation.

**Figure 4.**
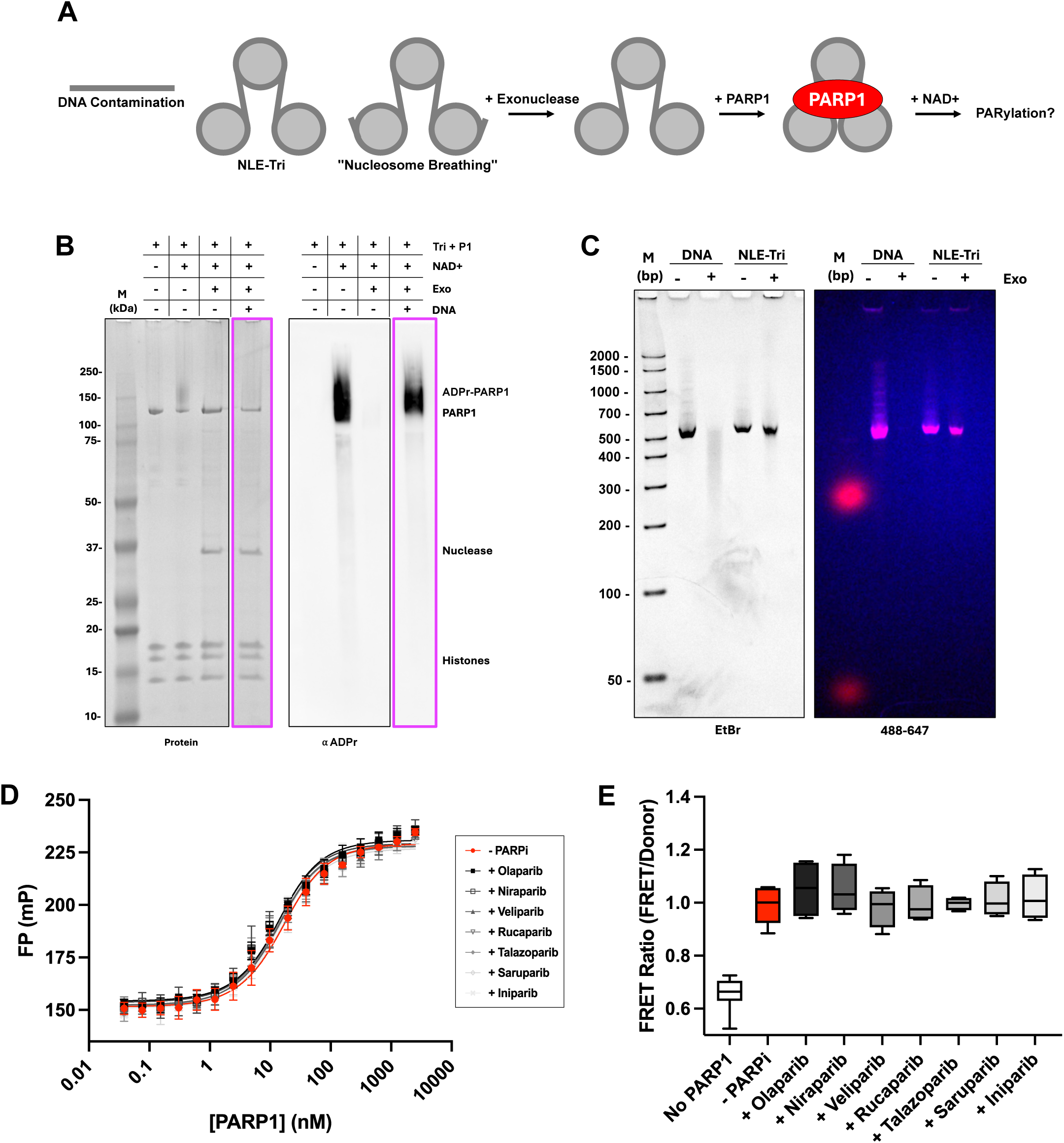
PARP1 is inactive in the presence of undamaged chromatin. (**A**) Experimental design. (**B**) PARP1-exonuclease activity protein gel (left) and ADP-ribose western blot (right). NLE-Tri, PARP1, and NAD+ are combined with or without exonuclease treatment and run on NuPAGE 4-12% Bis-Tris protein gels. Gels are either stained for protein or transferred and probed with anti-ADP-ribose antibody. Exonuclease activity was quenched and dsDNA was added back (magenta). (**C**) Analysis of exonuclease treated DNA and NLE-Tri. DNA and NLE-Tri were pretreated with exonuclease, then treated with Proteinase K. The resulting DNA fragments were purified over a spin column, analyzed by 6% DNA PAGE, and imaged for DNA (left) and 488/647 fluorescence (right). (**D**) NLE-Tri-PARP1 fluorescence polarization curve fit to the quadratic equation in the presence or absence of different PARP inhibitors (10 µM PARPi) at 200 mM KCl, n=3. (**E**) FRET-based chromatin compaction assay of PARP1-NLE-Tri in the presence or absence of PARPi as indicated. Replicates are done at 20 nM NLE-Tri, 50 nM PARP1, and 100 nM PARPi. Statistical significance was tested using the Brown-Forsythe and Welch ANOVA with Dunnett’s T3 multiple comparisons test, with no significance noted, n=4.

PARPi are used in targeted cancer therapy to effectively kill cancer cells lacking proficient DNA repair pathways (*24–28*). Currently, there are four FDA-approved inhibitors (Olaparib, Niraparib, Rucaparib, and Talazoparib) with several more in clinical trials (Veliparib, Saruparib, etc). Despite their clinical success, how PARPi affect PARP1 functions outside of DNA damage is understudied. Non-specific inhibition of PARP1 molecules could disrupt alternate PARP1 functions which could be detrimental to healthy cells. Thus, we wanted to determine if current approved and in-trial inhibitors could disrupt or alter the PARP1-chromatin architectural binding mode. To do this, we tested several types of inhibitors (Olaparib, Niraparib, Veliparib, Rucaparib, Talazoparib, Saruparib) in both the fluorescence polarization binding assay and FRET-based chromatin compaction assay. All affinities in the presence of inhibitors where identical to those measured in the absence of PARPi (**Fig. 4D**, **Table 3**), suggesting that these PARPi do not affect PARP1-NLE-Tri binding. We also tested Iniparib, a molecule originally thought to be a PARPi but later invalidated (*59*), as a negative control and found no effect. We also saw no disruption of FRET ratio compared to (-)PARPi or Iniparib, suggesting that these PARPi do not affect PARP1-induced chromatin compaction (**Fig. 4E**). Together, these results indicate that PARPi do not disrupt the PARP1 chromatin compacting binding mode. These findings are consistent with our result that PARP1 is enzymatically inactive when bound to undamaged chromatin. However, if PARP1 activity were required for downstream transcriptional regulation, it is likely that PARPi would disrupt this. More studies are needed to determine the effects of PARPi on other functions of PARP1, specifically ones in which PARP1 is active.

**Table 3.** PARP1 binding affinities in the presence of PARPi. Reported apparent dissociation constants (Kd_app_) for full-length PARP1 in the presence and absence of PARPi bound to NLE-trinucleosomes at 200 mM KCl. The affinities obtained were determined from the quadratic equation. R^2^ and replicate values are reported.

| Construct | Undamaged Chromatin (NLE-Tri) |  |  |
| --- | --- | --- | --- |
| | $K_{d_{app}}$ | $R^2$ | Replicates |
| - PARPi | $12.0 \pm 4.2$ | 0.980 | n=3 |
| + Olaparib | $9.1 \pm 1.9$ | 0.984 | n=3 |
| + Niraparib | $8.3 \pm 1.8$ | 0.968 | n=3 |
| + Veliparib | $9.2 \pm 3.2$ | 0.982 | n=3 |
| + Rucaparib | $9.0 \pm 2.2$ | 0.978 | n=3 |
| + Talazoparib | $7.9 \pm 1.2$ | 0.974 | n=3 |
| + Saruparib | $10.5 \pm 2.6$ | 0.981 | n=3 |
| + Iniparib | $9.7 \pm 5.7$ | 0.976 | n=3 |

## DISCUSSION

PARP1 is a highly abundant nuclear enzyme that dynamically associates with chromatin through both transient and stable binding (*60*). Here we show through rigorous biochemical and biophysical in vitro analysis that PARP1 exhibits a distinct chromatin binding mode that does not lead to activation. Characterization of full-length PARP1 bound to a minimalist model of undamaged chromatin (NLE-Tri: three nucleosomes connected by 22 bp DNA linker, without any accessible DNA ends) validates that one PARP1 molecule binds to two chromatin linkers simultaneously (**Fig.1**). This interaction brings the three nucleosomes closer together, a hallmark of chromatin compaction (**Fig. 2**). Characterization of various constructs of PARP1 where individual DNA binding domains were deleted shows that the Zn1, Zn2, Zn3, and BRCT domains all contribute to the PARP1-chromatin architectural binding mode (**Fig. 3**). These results are consistent with previous HDX-MS data that revealed protection of these same domains when bound to “intact” plasmid DNA (*41*). Interestingly, both the Zn2 and BRCT domains in the HDX-MS experiment showed almost full protection when bound to plasmid DNA as opposed to short DNA fragments, which would suggest their importance in this binding mode. We also saw the most significant disruptions in both binding and compaction of NLE-Tri when either the Zn2 or BRCT domains were deleted, confirming their contribution in this more physiologically relevant context of undamaged chromatin. Additionally, our data reveal that PARP1 is inactive when bound to undamaged chromatin, something that has not been shown before due to complications with DNA contamination (**Fig. 4**). With the engagement of the BRCT domain and the lack of engagement of the WGR domain, the cascade of conformational changes to displace the HD does not occur in this binding mode, and thus PARP1 remains inactive. Finally, since PARP1 is inactive when bound to NLE-Tri, PARPi do not affect disrupt this interaction under our conditions (**Fig. 4**). Altogether, we have found that the Zn1, Zn2, Zn3, and BRCT domains contribute to the binding and compaction of undamaged chromatin by engaging the DNA linkers in confirmation that does not lead to catalytic activation.

As an integral component of several DNA damage repair pathways, PARP1 is thought to constantly scan the genome for DNA lesions through intrastrand transfer (i.e. the “monkey bar” mechanism) (*41*, *56*). With evidence to suggest that ∼10^6^ PARP1 molecules are present in the nucleus (*61*), PARP1 activation via binding undamaged chromatin would be detrimental to the cell, resulting in depletion of NAD+, erroneous recruitment of the DNA damage repair machinery and subsequently, cell death. Although various studies have shown that PARP1 is activated by non-linker ended mono-nucleosomes and nucleosomal arrays, we have demonstrated that minute amounts of DNA contamination, or the transient dissociation of the ends of nucleosomal DNA obscure these results (*62*, *63*). Using exonuclease to control for this potential contamination has revealed that PARP1 is completely inactive in this context, countering previous results reported from our lab and others (**Fig. 4B**). These results are further validated by the lack of WGR engagement in the undamaged chromatin binding mode, as well as by the importance of Zn2 (**Fig. 3B&E**), which was shown to be dispensable for PARP1 activation (*15*, *43*). These findings highlight the importance of rigorously controlling in vitro PARP1 activity assays, as even trace DNA contamination is sufficient to activate PARP1 and confound interpretation of results.

From our data, we propose that the Zn1, Zn2, Zn3, and BRCT domains are essential for navigating the genome by binding chromatin linkers without affecting the HD domain that covers the active site (**Fig. 5A: 1**). We hypothesize that upon detecting a DNA lesion, a switch in domain engagement occurs in which the WGR displaces the BRCT domain. Upon WGR engagement, the HD undergoes a conformational change, allowing access to the catalytic binding pocket (**Fig. 5A: 2**). In this activated state, NAD+ binds, and PARP1 can PARylate itself (**Fig. 5A: 2**), as well as histones in the presence of HPF1 (**Fig. 5A: 3**). Histone PARylation serves to destabilize the nucleosome and relax chromatin to better allow access to the damage site (**Fig. 5A: 3**). Further, PARylation is essential for signaling the presence of damage and subsequently directing the DNA damage repair machinery to the damage site (**Fig. 5A: 4**). Finally, PARG and other glycohydrolases restore PARP1 and recycle PAR. Thus, PARP1 is no longer activated and can continue scanning the genome or other lesions.

**Figure 5.**
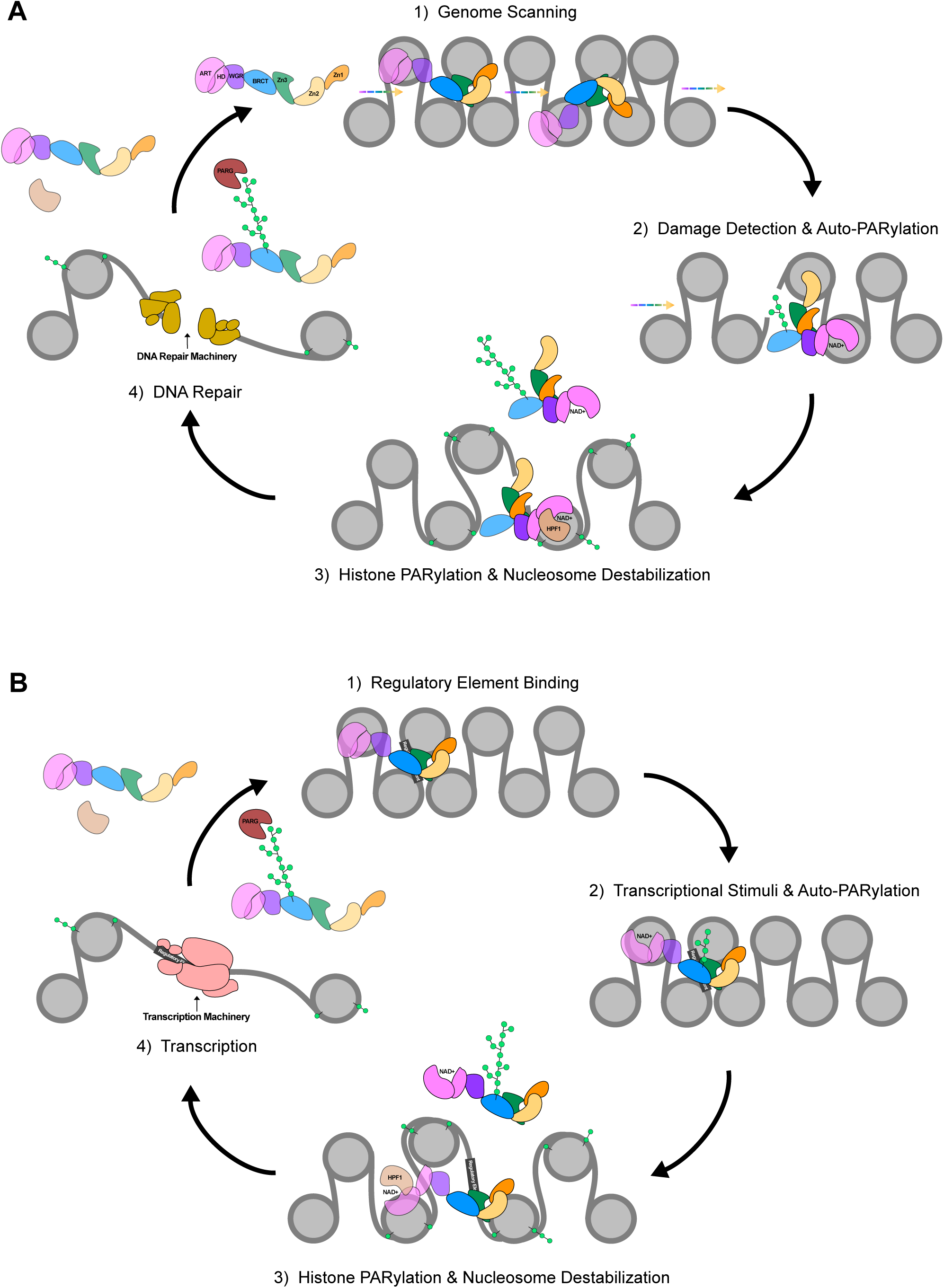
Model for the role of PARP1-inactive chromatin binding mode in DNA repair and transcription. **(A)** DNA repair: 1) PARP1 searches the genome for DNA damage by simultaneously binding two chromatin linkers via Zn1, Zn2, Zn3, and BRCT. 2) Upon detecting damage, the WGR engages and the BRCT is displaced from DNA. WGR engagement leads to HD unfolding, which in turn enables access to the NAD+ binding pocket. NAD+ binds and PARP1 auto-PARylates itself. 3) Auto-PARylated PARP1 dissociates from the DNA. Activated PARP1 forms a composite active site with HPF1, switching PARylation to histones. Histone PARylation destabilizes the nucleosome. 4) The relaxed chromatin state enables the DNA repair machinery to readily access and repair the damaged DNA. PARG and other glycohydrolases degrade PAR, returning PARP1 to the non-active state to continue scanning the genome for damage. **(B)** Transcription: 1) PARP1 stably binds regulatory DNA elements and 2) becomes activated upon a yet unidentified transcriptional stimulus, freeing access to the NAD+ binding pocket and leading to PARP1 auto-PARylation. 3) Auto-PARylated PARP1 dissociates from the DNA. If HPF1 is present during transcription, activated PARP1 can form a composite active site with HPF1, targeting PARylation to histones. Histone PARylation could then destabilize the nucleosome. 4) The relaxed chromatin state enables access to the transcriptional machinery. PARG and other glycohydrolases degrade PAR, returning PARP1 to the non-active state to rebind regulatory element to control transcription.

PARP1 also plays a role in regulating genome architecture and subsequently, gene expression, but contradicting mechanisms have been described. For example, PARP1 bound to chromatin represses RNA Pol II, yet PARP1 also promotes the formation of open chromatin at promoters of highly expressed genes (*36*, *42*, *64*). In *Drosophila*, PARP1 is located at the *Hsp70* gene and is activated in response to heat shock. Upon activation, PARP1 redistributes along *Hsp70* to PARylate and displace histones, allowing efficient transcription by RNA pol II (*65–67*). In this context, PARP1 is both bound to a specific gene and essential for transcriptional activation. As previously shown, a population of PARP1 molecules stably bind chromatin in live human cells (*60*). ChIP-seq studies in human cells have shown that PARP1 binds regulatory elements near transcriptional start sites (*37*, *38*, *68*). These interactions suggest a similar mechanism of transcriptional regulation as seen in *Drosophila*. In this context, we propose that PARP1 binds the chromatin linkers using the Zn1, Zn2, Zn3, and BRCT domains as a way to regulate gene expression (**Fig. 5B**). Here, the Zn1, Zn2, Zn3, and BRCT domains engage the regulatory element and repress RNA pol II binding through binding/compaction (**Fig 5B: 1**). Upon unspecified and as yet uncharacterized stimuli (transcription factor, epigenetic marks, hormones, etc.), PARP1 activates and PARylates itself (**Fig 5B: 2**) and potentially histones, if HPF1 is present (**Fig 5B: 3**). Histone PARylation would contribute to nucleosome destabilization and chromatin relaxation to better allow the transcriptional machinery to access the site. PAR chains could also act as a signal for transcriptional activation, recruiting transcription machinery to the start site. Finally, PARG and other glycohydrolases can come to restore PARP1 and recycle PAR, thus allowing PARP1 to rebind/repress the regulatory element. This model is more speculative as it is unknown whether HPF1 and histone PARylation play a role in transcription. More studies are needed to further probe the role of PARP1 and ADP-ribosylation in the context of gene expression and regulation.

PARP1 is dynamically associated with chromatin through a wide variety of processes and mechanisms. While distinguishing these processes and mechanisms into distinct functions has been difficult, we have determined how PARP1 directly engages undamaged chromatin. In this study, we found that PARP1 binds the chromatin linker with high affinity, resulting in compaction of the nucleosomal array. This interaction occurs through the engagement of the Zn1, Zn2, Zn3, and BRCT domains but does not lead to PARP1 enzymatic activation. We additionally report that PARPi do not disrupt PARP1-chromatin binding or compaction. Our proposed model clarifies previously contradicting ideas by showing how PARP1 may use this inactive compaction state to bind chromatin until its activation in their DNA damage or transcription context.

## MATERIALS AND METHODS

### Protein expression and purification

Full-length PARP1 was expressed in SF9 insect cells with Baculovirus purchased from Kinakeet Biotech. SF9 cells were thawed from - 80°C, resuspended in 27°C warmed media (ESF 921 – Expression Systems), and spun down for 5 minutes, 900 rcf at room temperature. The pellet was resuspended with media, transferred to a 125 mL culture flask and incubated while shaking at 27°C, 130 rpm. Cells were subcultured to a final 3 L culture flask at 1.8 x 10^6^ total cells/mL and inoculated with 12 mL of virus (titer: >1x 10^8^) for 3-4 days. Cells were pelleted for 20 minutes at 3000 rpm, flash frozen and stored at-80°C.

Cell pellets were thawed from-80°C and resuspended on ice in lysis buffer (25 mM Tris-HCl pH 7.5, 500 mM NaCl, 30 mM Imidazole pH 7.5, 0.5 mM TCEP, 1 mM AEBSF, 0.1% NP40, 1 mM PMSF, 0.5 μg/mL Leupeptin, 0.7 μg/mL Pepstatin, 0.5 μg/mL Antipain, 0.5 μg/mL Aprotinin, 1 mM Benzadmidine, 2 EDTA-free ULTRA protease inhibitor tablets). Resuspended pellets were stirred on ice with 1.5 mM MgCl_2_ and 3 μL benzonase/liter of cells for 5 minutes then sonicated for 4 minutes “on” recall 3 at 30%. Cell lysates were centrifuged for 25 minutes, 18,000 rpm at 4°C and the pellet was discarded. The supernatant was filtered using a 0.45μm filter and loaded onto a HisTrap HP column and eluted with a linear gradient (0-100% B buffer) of Ni B buffer (25 mM Tris-HCl pH 7.5, 500 mM NaCl, 1 M Imidazole pH 7.5, 0.5 mM TCEP, 1 mM AEBSF). Further purification included a HiTrap Heparin eluted in Heparin High Salt Buffer (50 mM Tris pH 7.5, 0.1 mM EDTA pH 8.0, 1 M NaCl, 0.1 mM TCEP, and 1 mM AEBSF) and S200 Size Exclusion column eluted in S200 Buffer (25 mM HEPES pH 7.0, 0.1 mM EDTA pH 8.0, 150 mM NaCl, 0.1 mM TCEP, 1 mM AEBSF).

PARP1 mutants (ΔZn1, ΔZn2, ΔZn3, ΔBRCT, ΔWGR, ΔCAT, N-term, and C-term) were purchased from the Histone Source (Fort Collins, CO). PARP1 mutants were expressed in E. coli and purified in the same manner as full-length PARP1 (HisTrap HP, HiTrap Heparin, and S200 Size Exclusion Chromatography).

### NLE-Trinucleosome (NLE-Tri) assembly and quality control

#### Dual-labeled 485 bp DNA amplification and purification

Dual-labeled trinucleosomes were assembled on 485 bp dsDNA containing the 607, 626, and 603-Widom sequences (*69*), separated by 22 bp of linker DNA (see sequences). The DNA sequence was analyzed using NuPoP to ensure optimal nucleosome positioning (*70*). PCR was employed to amplify large quantities of dual-labeled DNA using IDT gBlock, 5’ attached AlexaFluor 488 forward primer, 5’ attached AlexaFluor 647 reverse primer (see sequences), and OneTaq 2X Master Mix. Amplified DNA was purified off a MonoQ 10/100 8 mL column in DNA elution buffer (10 mM HEPES pH 7.5, 1 mM EDTA pH 8.0, 1M NaCl) using a linear gradient, concentrated by ethanol precipitation, and resuspended in molecular grade water. DNA was analyzed on an Invitrogen 6% DNA retardation gel at RT for 1 hour at 100V and imaged for fluorescence at 488 nm and 647 nm channels. Gels were then stained with ethidium bromide (EtBr) and imaged for DNA.

#### Histone octamer reconstitution and purification

Human histone octamer reconstitution and purification were performed as previously described (*71*). Briefly, lyophilized human histone proteins (H2A, H2B, H3, and H4) were purchased from the Histone Source. Equimolar amounts of each histone were combined in unfolding buffer (6 M Guanidinium HCl, 20 mM HEPES pH 7.0, 5 mM DTT) and dialyzed into refolding buffer (2 M NaCl, 10 mM HEPES pH 7.0, 1m M EDTA, 5 mM β-mercaptoethanol) overnight. Octamer was purified using Superdex S200 (16/60) 120 mL column. Fractions were run on an Invitrogen precast 4-12% Bis-Tris gel. Fractions with the correct ratio of histones were concentrated and flash frozen in glycerol salt buffer (2 M NaCl, 50% glycerol) at a final concentration of 10% glycerol.

#### Dual-labeled trinucleosome reconstitution

Dual-labeled trinucleosomes were first assembled by combining different ratios of histone octamer to dual-labeled DNA. Trinucleosomes were then dialyzed in a continuous salt gradient overnight; followed by an additional overnight dialysis in 0M refolding buffer (20 mM MES pH 6.0, 1 mM EDTA, 0.1 mM TCEP) (*72*). Samples were digested using SacI/XhoI for 2 hours at 37°C and analyzed by 5% native PAGE in TBE at 4°C for 60 minutes at 150 V. Gels were imaged for fluorescence at 488 nm and 647 nm channels. Gels were then stained with EtBr and Blazin’ Blue and imaged for DNA and protein, respectively. Saturated trinucleosomes (i.e. trinucleosomes with no free DNA and no undigested trinucleosome) were further tested for purity by mass photometry and negative stain electron microscopy. Pure saturated trinucleosomes were then scaled up for downstream assays.

#### Unlabeled trinucleosome reconstitution

Unlabeled trinucleosomes were reconstituted on 485 bp dsDNA containing three 601-Widom sequences (*69*) separated by 22bp linker DNA. The DNA sequence was cloned and purified to obtain large quantities as previously described (*72*). Unlabeled trinucleosomes were assembled as previously mentioned and tested for purity using BamHI restriction digest and mass photometry. Pure saturated trinucleosomes were then scaled up for downstream assays.

#### Fluorescence polarization (FP) binding assay

For trinucleosome binding experiments, 5 µM PARP1 was serially diluted 1:2 into a 384 well plate containing 10 µL of binding buffer (50 mM HEPES pH 7.0, 100 mM or 200 mM KCl, and 0.01% NP40). Dual-labeled trinucleosomes were added at a final concentration of 10 nM. Plates were incubated at RT for 1 minute. PARP1 domain deletions FP assays were performed identically. PARPi FP assays were performed in the presence of 10 µM inhibitor with the 200 mM KCl binding buffer.

For DNA binding experiments, 1µM PARP1 was serially diluted 1:2 into a 384 well plate containing 10 µL of binding buffer. p18mer was added at a final concentration of 5 nM. Plates were incubated at RT for 1 minute. PARP1 domain deletions FP assays were performed identically.

For each experiment, FP (mP) was measured using a BMG Labtech CLARIOstar plate reader with excitation at 488nm. The FP baseline was set using labeled trinucleosome or labeled DNA only controls to establish the unbound baseline. All trinucleosome FP experiments were fit to the quadratic equation (1), where Ymin and Ymax are the minimum and maximum FP signals, respectively, Kd is the experimental binding constant, L is the trinucleosome concentration, and X is the PARP1 concentration.

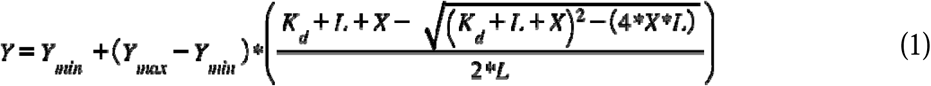

All DNA FP experiments were fit to the 4-parameter binding curve (2), where Bottom and Top are the minimum and maximum FP signals, respectively, EC50 is the experimental binding constant, X is the PARP1 concentration, and Hill Slope is the hill coefficient.

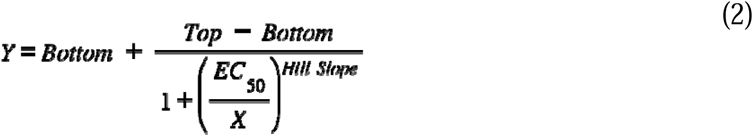

Replicate experimental binding constant values were plotted in a bar graph to obtain an average Kd_app_ and standard deviations. All FP experiments were repeated more than three times from independent dilution series prepared on different days.

#### Electrophoretic mobility shift assay (EMSA)

Unlabeled trinucleosome (50 nM) was mixed with 0X-4.5X molar equivalents of FL PARP1 in binding buffer (50 mM HEPES pH 7.0, 100 mM KCl). Samples were incubated at RT for 1 minute, loaded into a 4% native TBE gel, and run at 4°C for 75 minutes at 150 V. EMSA gels were first stained in EtBr and imaged for DNA, then stained in Blazin’ Blue and imaged for protein. PARP1 domain deletion EMSA gels were tested at different molar equivalents of protein: ΔZn1: 0-4X, ΔZn2: 0-7X, ΔZn3: 0-4.5X, ΔBRCT: 0-23X, ΔWGR: 0-4.5X, ΔCAT: 0-17X.

#### Mass photometry (MP)

Mass photometry samples were measured on the Refeyn TwoMP mass photometer. Glass slides were cleaned with isopropanol and deionized water and dried with N_2_ gas. All samples were prepared with 50 nM trinucleosome and 0-3X PARP1 in binding buffer and diluted to ∼15nM in the same buffer. 60 second videos were recorded using Refeyn AcquireMP. ß-amylase and Thyroglobin was used as mass standards to convert contrast signal to molecular weight. All movies were analyzed in DiscoverMP.

#### Micrococcal nuclease (MNase) digestion

Unlabeled trinucleosome (200 nM) in the presence (400 nM) and absence of PARP1were digested with increasing units of MNase (0 U, 6.25 U, 12.5 U, 25 U), 1X MNase buffer, and 0.1 mg/mL BSA. Reactions were quenched with 0.1 M EDTA and digested with 20 µg of ProteinaseK at 50°C for 30 minutes. DNA was purified using the Qiagen MinElute kit and run on an Invitrogen 6% DNA retardation gel at 15 ng/µL at RT for 1 hour at 100 V. The 485 bp band intensity was analyzed in FIJI and normalized to the corresponding control intensity (0 U) to get the % Protection. Unpaired t-test with Bonferroni-Dunn’s multiple comparisons test was used to assess statistical significance n=3. N-term PARP1 MNase digest was performed identically.

#### Förster resonance energy transfer (FRET) compaction assay

For PARP1 and PARP1 domain deletion assays, 20 nM dual-labeled trinucleosome was mixed with 0X-5.5X molar equivalents of PARP1 at a final volume of 20 µL in the 100 mM KCl FP binding buffer. FRET intensities were measured using a BMG Labtech CLARIOstar plate reader with excitation at 488 nm and emission at 647 nm. 2.5X molar equivalents of PARP1 (and PARP1 domain deletions) was used to compare to no PARP1 after determining little to no aggregation present. Raw FRET intensities were normalized to raw donor (488 nm) intensities to get the FRET ratio. FRET ratio values were plotted in Graphpad Prism in a box and whisker plot with n=5 replicates. Brown-Forsythe and Welch ANOVA with Dunnett’s T3 multiple comparison adjustment were used to determine statistical significance. PARPi FRET assays were done in the presence of 2X inhibitor (∼100 nM).

#### Negative stain electron microscopy

Negative stain samples were prepared by mixing 20 nM unlabeled trinucleosome alone or with 2.5X PARP1 (50 nM) in binding buffer. 4 µL of sample was applied to a glow discharged 400-mesh copper grid (Electron Microscopy Services) with a continuous carbon-support layer. After 30 seconds on incubation, the grids were stained with 2% (w/v) uranyl formate solution in five 35 µL drops and blotted to dry. Grids were screened using the FEI Tecnai T12 transmission electron microscope (TEM) and data was collected using the FEI Tecnai F20 TEM. Data was processed in cryoSPARC to create ab-inito reconstructions.

#### Western blot

Exonuclease treated trinucleosomes were digested using 100 nM trinucleosome, 3 units T5 exonuclease, 1X NEB buffer, 3 units of Mung Bean Nuclease, and 1X Mung Bean Nuclease buffer and incubated for 30 minutes at 30°C in binding buffer. Trinucleosomes without exonuclease were also incubated for 30 minutes at 30°C in binding buffer with 1X NEB buffer and 1X Mung Bean Nuclease buffer. Following incubation 100 nM PARP1 was added to each reaction. 100 µM NAD+ was added to all reactions except for the negative control. Reactions in which dsDNA was later added were first quenched by adding 45 mM EDTA pH 8.0 and after, 200 nM NLE-Tri DNA (485 bp). 2X Laemmli was added to each reaction before running on two Invitrogen precast 4-12% Bis-Tris gel for 200 V at room temperature for 30 minutes. One gel was stained with Blazin’ Blue for protein imagine while the other gel was transferred to a PDVF membrane using the Invitrogen Power Blotter System for 7 minutes at 25 V. Following transfer, the membrane was placed in blocking buffer (5% milk powder in 1X TBST (20 mM Tris-HCl pH 7.6, 150 mM NaCl, 0.1% Tween 20)) for 1 hour. After 1 hour, blocking buffer was replaced with primary buffer (3% milk powder in 1X TBST, with 2 µL Cell Signaling Technology Poly/Mono ADPr Rabbit mAb (89190S)) for 1 hour. After 1 hour, primary was rinsed off the membrane with 1X TBST 3-4x. Secondary buffer (3% milk powder in 1X TBST, with 2 µL Cytiva Anti Rabbit HRP (45-000-682)) was then added for 1 hour. After 1 hour, secondary was rinsed off the membrane with 1X TBST 3-4x. 1:1 stable peroxide and luminol/enhancer was added to the membrane for 1-2 minutes and imaged using chemi and red ladder.

#### Sequences

Dual-labeled NLE-Tri gBlock (Widom 607, 626, and 603 are underlined respectively): GAGGCCTAACGAGCGCCTTGGTATCCGGAACGGGTTCTAGTACCCGTTAGCGTGGTTTAGAGGGGCAAAGGAACATCTTTCCCCCCCCCGAGATACGGGCACCGTTCGGACCCTGGTTAGTCCAGTGCTACTGCCGGTTCCTAGCCCTTCGAGCTCTCACAGATCTGAAAAGCCTTGTCGCTCTGCGATTCGATGAGAGTCGACCGGTGCCGGGTATTGTATCGCCGCTTGACTCTGGAATGGCGTCTACGTAGCGCTAAGCACTATTAGAGTCCTCTCCTGCATCTCCAAGGATAGTGCCTATAAATCGTCCACCTTCCTCGAGTCACAGCTCTGAAGCCGTAAAATAATCGACACTCTCGGGTGCCCAGTTCGCGCGCCCACCTACCGTGTGAAGTCGTCACTCGGGCTTCTAAGTACGCTTAGCGCACGGTAGAGCGCAATCCAAGGCTAACCACCGTGCATCGATGTTGAAAGAGGCCCTC

Fwd Primer: 5’Alex488N-GAGGCCTAACGAGCGC

Rev Primer: 5’Alex647N-GAGGGCCTCTTTCAACATCG

Unlabeled NLE-Tri (Widom 601 sequences are underlined): ATCTGAGAATCCGGTGCCGAGGCCGCTCAATTGGTCGTAGACAGCTCTAGCACCGCTTAAACGCACGTACGCGCTGTCCCCCGCGTTTTAACCGCCAAGGGGATTACTCCCTAGTCTCCAGGCACGTGTCAGATATATACATCCGATTTCGGATCCTCACAGATCTGAAATCTGAGAATCCGGTGCCGAGGCCGCTCAATTGGTCGTAGACAGCTCTAGCACCGCTTAAACGCACGTACGCGCTGTCCCCCGCGTTTTAACCGCCAAGGGGATTACTCCCTAGTCTCCAGGCACGTGTCAGATATATACATCCGATTTCGGATCCTCACAGATCTGAAATCTGAGAATCCGGTGCCGAGGCCGCTCAATTGGTCGTAGACAGCTCTAGCACCGCTTAAACGCACGTACGCGCTGTCCCCCGCGTTTTAACCGCCAAGGGGATTACTCCCTAGTCTCCAGGCACGTGTCAGATATATACATCCGAT

## Supporting information

fiorenza et al, supplementary data

## Acknowledgments

We thank Dr. Johannes Rudolph and Dr. Annette Erbse for valuable discussion and technical help, and Dr. Nicole Hoitsma for critical reading of the manuscript.

## Funding

National Institute of Health Molecular Biophysics Training Grant T32 GM145437 (ACF)

National Cancer Institute RO1 CA218255 (KL)

Howard Huges Medical Institute (KL)

Shared Instrument Pool Core Facility RRID: SCR_018992

Boulder Electron Microscopy Core Facility RRID: SCR_001432

## Author contributions

Conceptualization: ACF, KL

Methodology: ACF

Investigation: ACF, MA

Data Curation: ACF Validation: ACF

Formal Analysis: ACF

Visualization: ACF

Supervision: KL

Project Administration: KL

Funding Acquisition: KL

Writing—original draft: ACF, KL

Writing—review & editing: ACF, MA, KL

## Data, code, and materials availability

All data needed to evaluate and reproduce the results in the paper are present in the paper and/or the Supplementary Materials. This study did not generate any new materials.”

