## Supplementary material for "PARP1 Exhibits an Enzymatically Inactive Chromatin Binding Mode": fiorenza et al, supplementary data

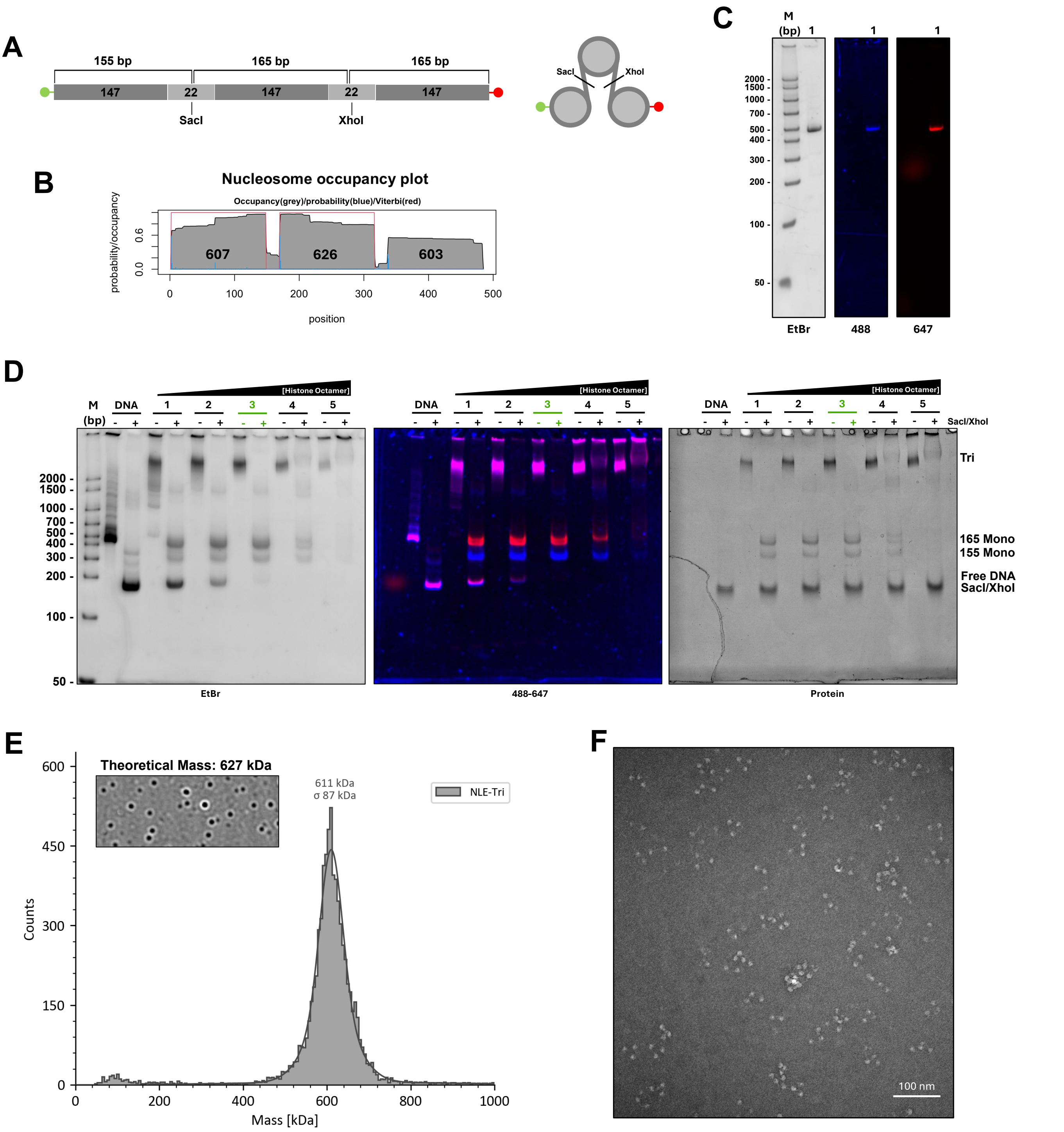


**Figure S1. Dual-labeled NLE-Tri-22 design, reconstitution, and quality control. (A)** Schematic of NLE-Tri-22 DNA and assembled trinucleosome. NLE-Tri-22 DNA contains three different Widom 147 bp nucleosome positioning sequences (607, 626, 603, respectively (65), see methods.) separated by 22 bp of internal linker DNA. Each linker harbors SacI or XhoI restriction sites for downstream quality control. AlexaFluor 488 and AlexaFluor 647 are 5' attached to opposite ends of the DNA. **(B)** Predicted nucleosome occupancy plot of NLE-Tri-22 DNA sequence from NuPoP (66). Plot shows the probability that a nucleosome covers a given position (grey), starts at a given position (blue), and the Vitberbi prediction (red). **(C)** PCR amplified dual-labeled NLE-Tri-22 DNA gel imaged for DNA (left), 488 nm (middle), and 647 nm (right). **(D)** SacI/XhoI restriction digest of NLE-Tri-22 titration. Different ratios of octamer are used to find the optimal degree of saturation (sample 3, green) where free DNA is absent, and linkers are fully digested. **(E)** Mass photometry analysis of optimal trinucleosome sample with a representative ratiometric contrast frame. Theoretical and observed masses are indicated. **(F)** Negative stain EM micrograph of optimal trinucleosome sample.

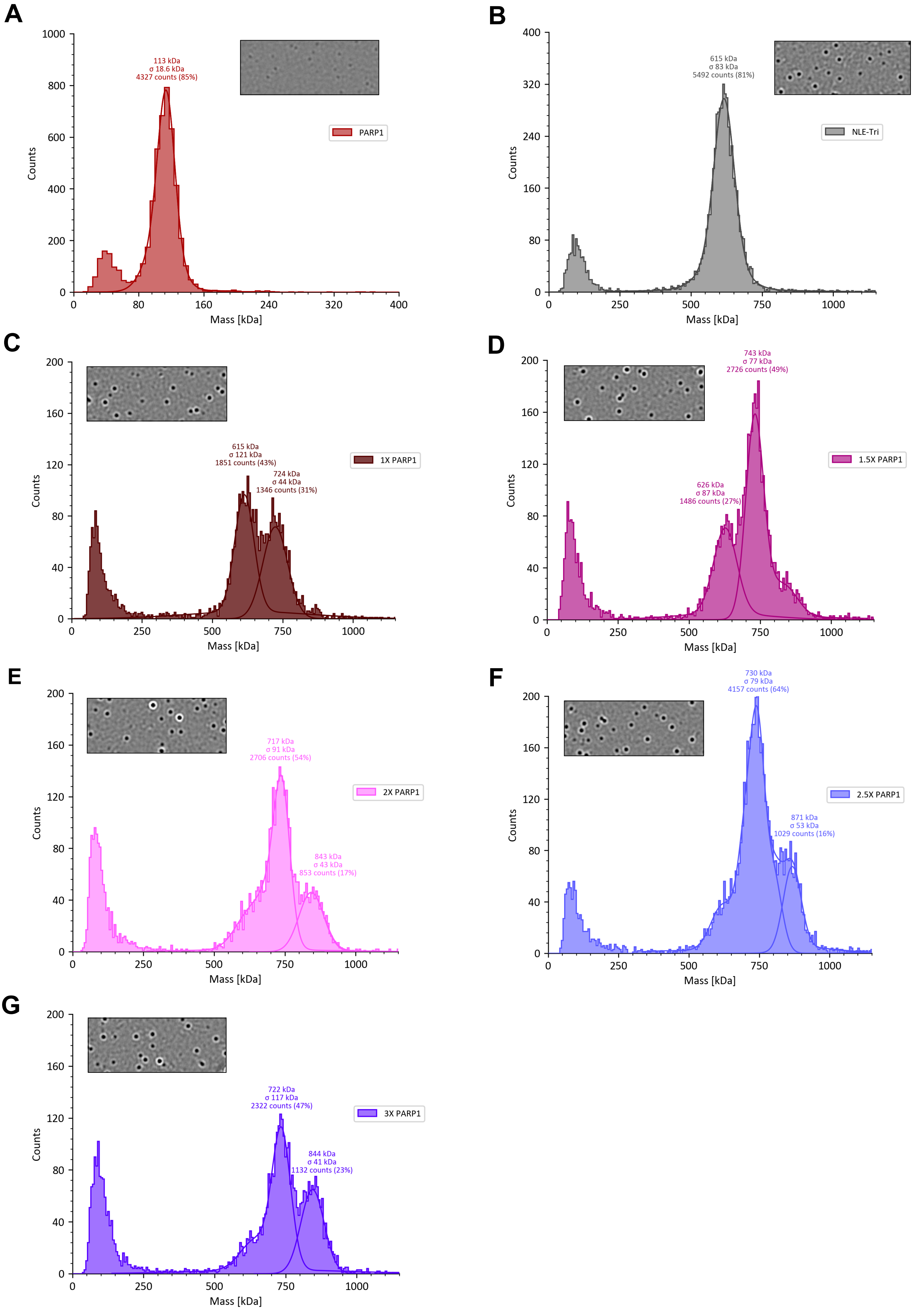


**Figure S2. Individual mass photometry analysis of PARP1 (A), NLE-Tri (B), and PARP1-NLE-Tri ratios of 1X (C), 1.5X (D), 2X (E), 2.5X (F), 3X (G).** Each mass photometry histogram is shown with a representative ratiometric contrast frame. Observed masses are indicated with sigma, number of counts, and percentage.

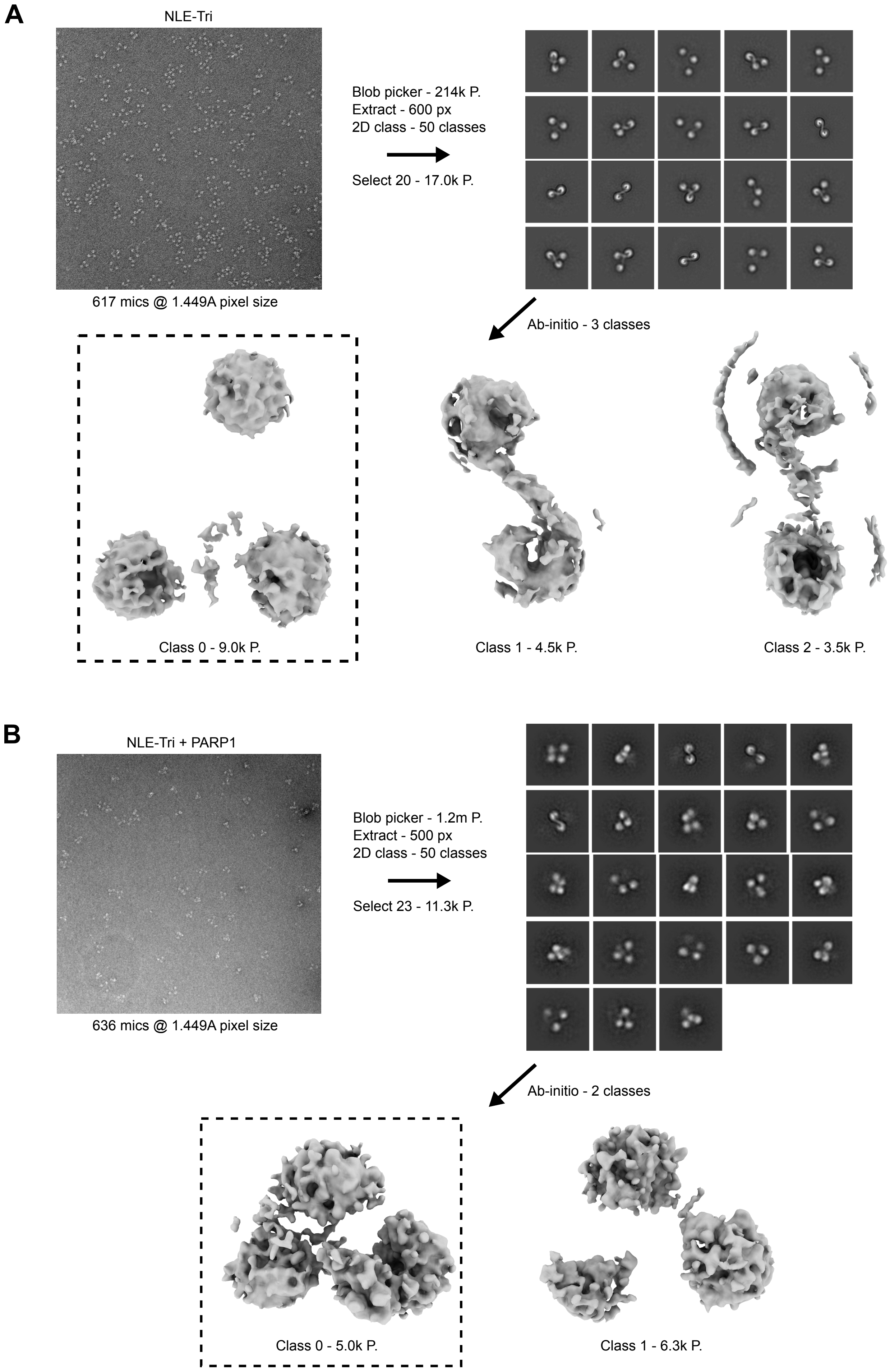


**Figure S3. Negative stain analysis workflow of (A) NLE-Tri alone; (B) NLE-Tri + PARP1.** All data analysis was done in CryoSPARC. Particles were picked using blob picker, inspected, extracted, and 2d classified. Following 2D classification, reasonable classes were selected and subjected to ab-inito reconstruction. One ab-inito class was selected for each dataset for analysis in ChimeraX.

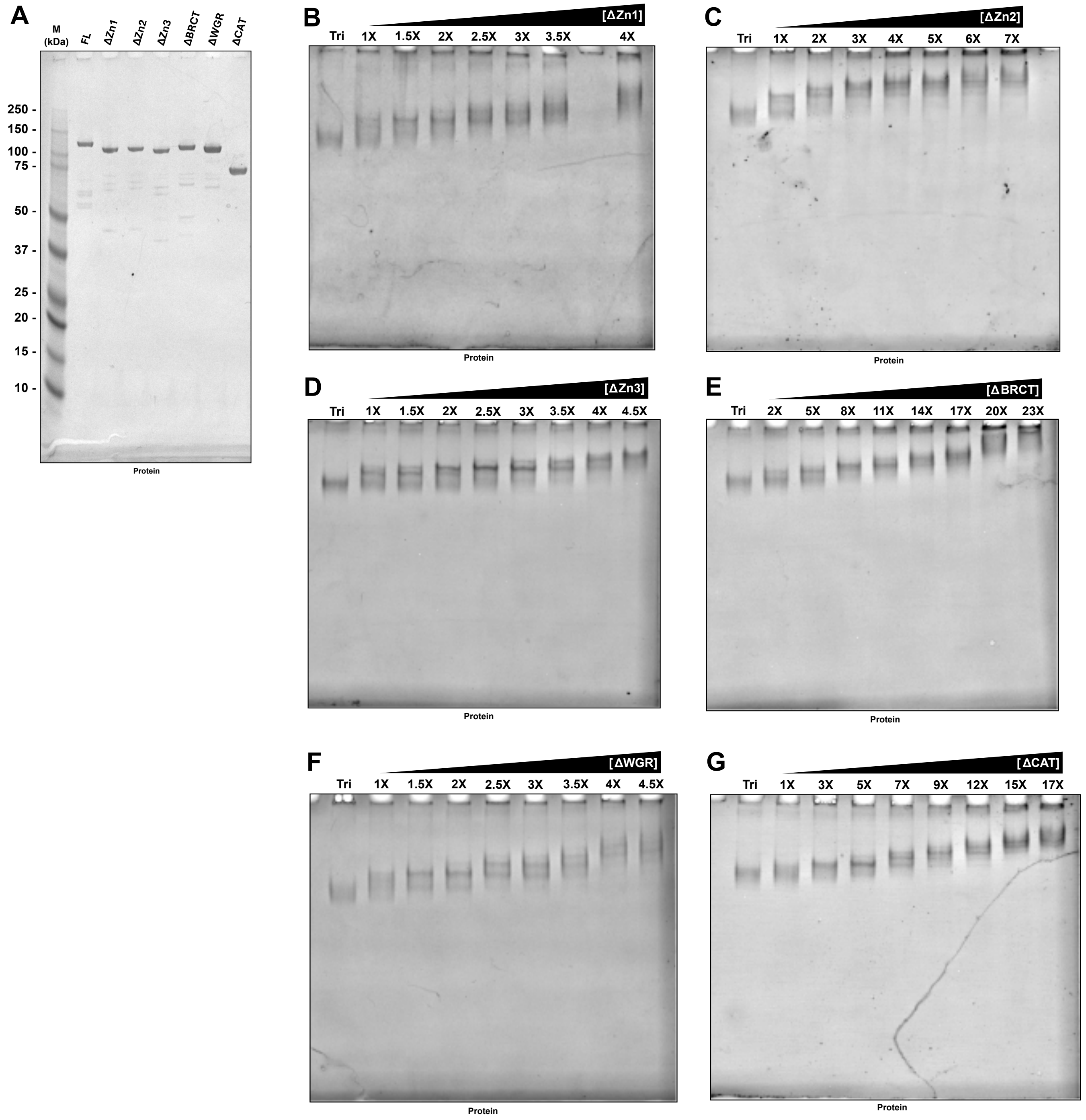


**Figure S4. PARP1 constructs with individual domain deletions retain the ability to bind NLE-Tri. (A)** SDS-PAGE showing purified full length (FL) PARP1 and each domain deletion construct used in this study (see Fig. 3A). EMSA of NLE-Tri-22 with **(B)** ΔZn1, **(C)** ΔZn2, **(D)** ΔZn3, **(E)** ΔBRCT, **(F)** ΔWGR, and **(G)** ΔCAT. All EMSAs were performed with indicated concentrations of PARP1 protein titrated into a fixed amount of NLE-Tri (50 nM). Protein molar equivalences are indicated above each lane.

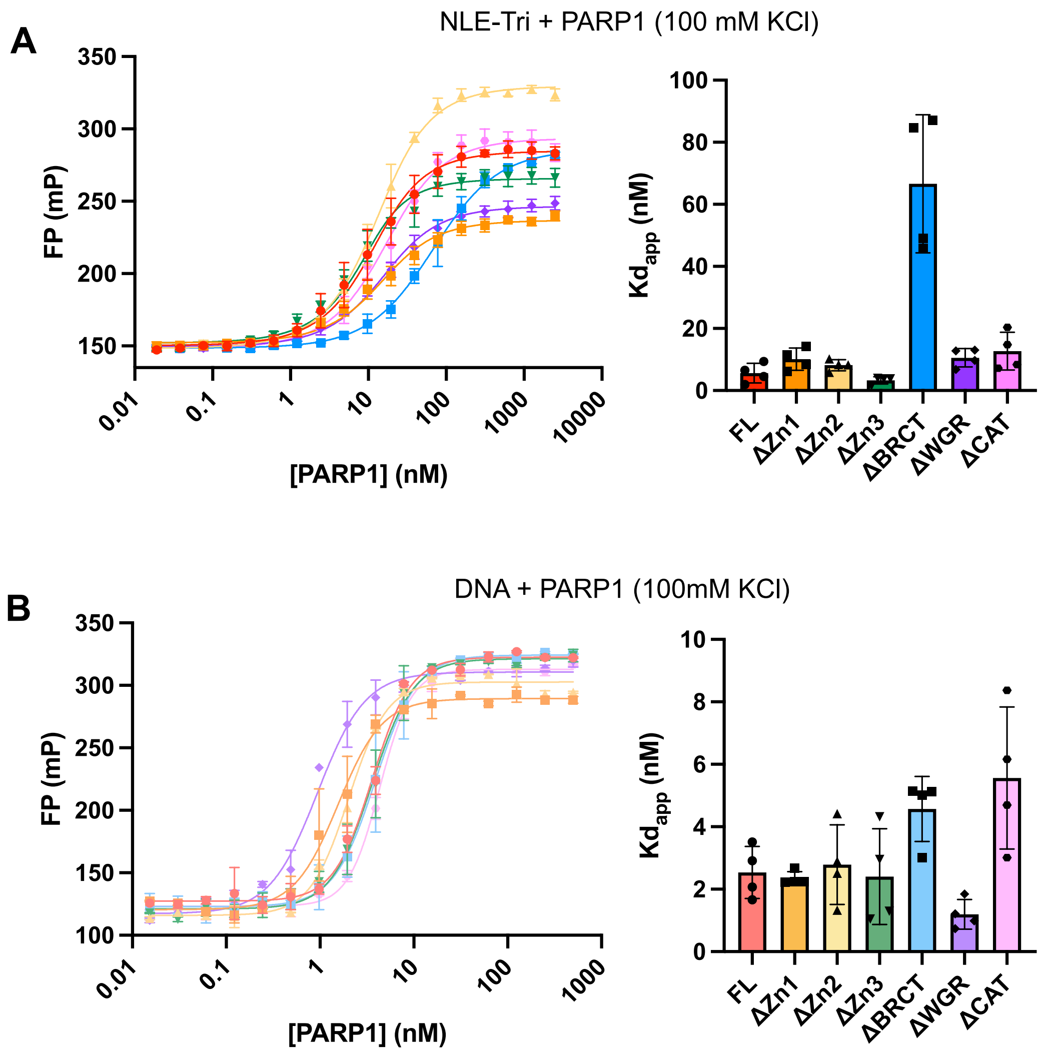


**Figure S5. PARP1 interaction with NLE-Tri and p18mer, measured by fluorescence polarization at 100 mM KCl. (A)** FP curve fit to the quadratic equation (left) and replicate affinities (right) of each construct bound to NLE-Tri at 100 mM KCl, n=4. **(B)** Fluorescence polarization curve fit to the four-parameter logistic curve (left) and replicate affinities (right) of each construct bound to p18mer at 100 mM KCl, n=4.

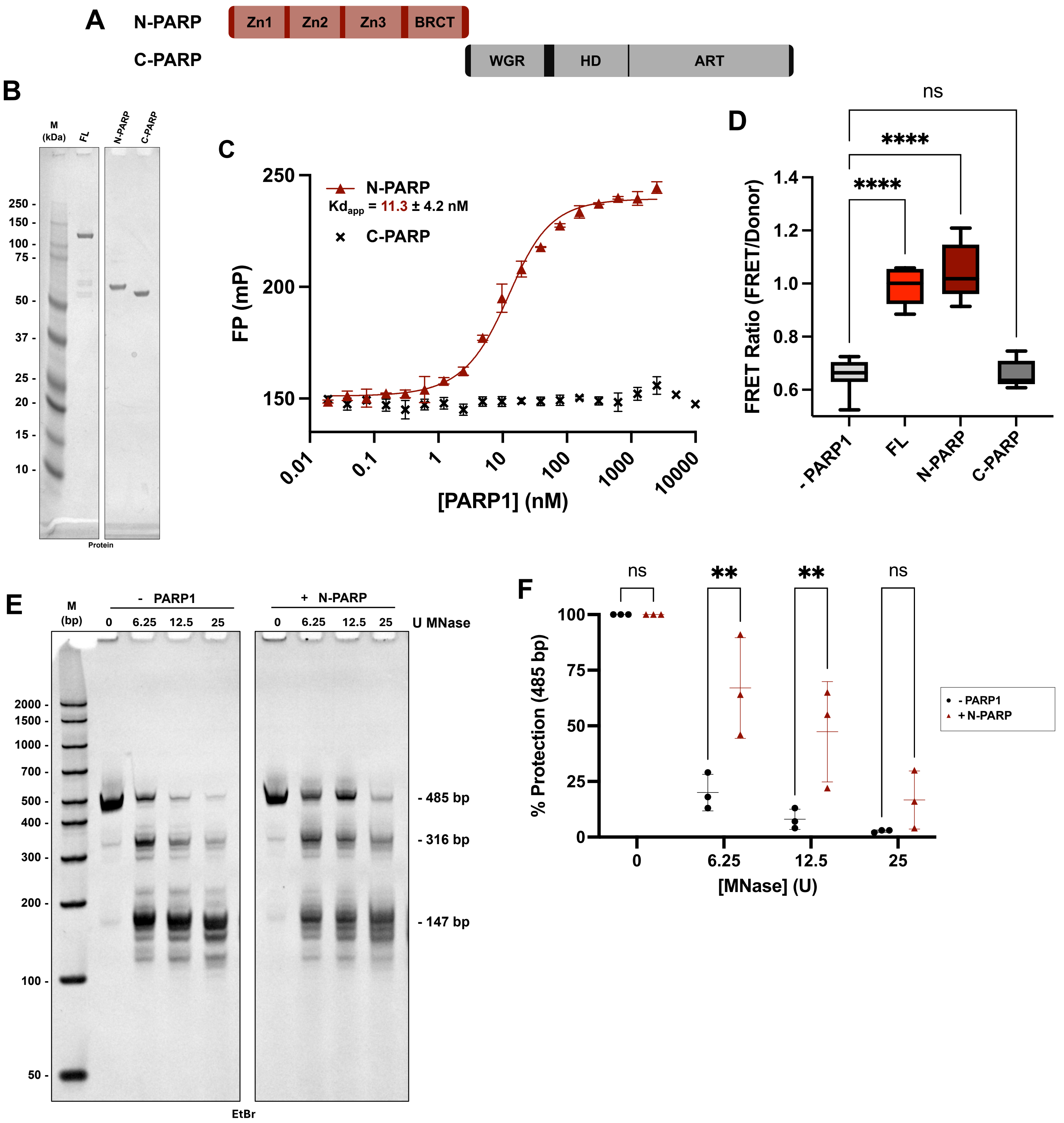


**Figure S6. N-PARP mimics full-length PARP1 in its chromatin architectural binding mode. (A) N-PARP and C-PARP domain map schematics. (B)** SDS PAGE showing purified full length (FL) PARP1, N-PARP, and C-PARP proteins. **(C)** N-PARP (maroon) and C-PARP (black X) fluorescence polarization at 100 mM KCl. N-PARP curve was fit to the quadratic equation. **(D)** FRET-based chromatin compaction assay replicates of FL, N-PARP, and C-PARP, compared to (-) PARP1. FRET ratio was determined by normalizing each FRET intensity to the 488 nm donor intensity. 50 nM protein was chosen for replicates, n=5. Statistical significance was determined using the ordinary one-way ANOVA with Dunnett's T3 multiple comparisons correction p<0.0001(****). **(E)** Representative MNase digest (analyzed by PAGE) in the presence and absence of N-PARP with increasing units of MNase, as indicated above each lane. **(F)** Quantitative analysis of the percent protection of the 485 bp band over increasing units of MNase. FIJI was used to determine each 485 bp band intensity. Each 485 bp band intensity was normalized to the corresponding 485 bp control (0 U) in the presence (maroon triangle) and absence (black circle) of N-PARP to determine percent protection. Statistical significance was determined using the unpaired t-test with Bonferroni-Dunn multiple comparison correction p<0.001 (***), n=3.
